# Reduced Default Mode Network Functional Connectivity in Patients with Recurrent Major Depressive Disorder

**DOI:** 10.1101/321745

**Authors:** Chao-Gan Yan, Xiao Chen, Le Li, Francisco Xavier Castellanos, Tong-Jian Bai, Qi-Jing Bo, Guan-Mao Chen, Ning-Xuan Chen, Wei Chen, Chang Cheng, Yu-Qi Cheng, Xi-Long Cui, Jia Duan, Yi-Ru Fang, Qi-Yong Gong, Wen-Bin Guo, Zheng-Hua Hou, Lan Hu, Li Kuang, Feng Li, Kai-Ming Li, Tao Li, Yan-Song Liu, Zhe-Ning Liu, Yi-Cheng Long, Qing-Hua Luo, Hua-Qing Meng, Dai-Hui Peng, Hai-Tang Qiu, Jiang Qiu, Yue-Di Shen, Yu-Shu Shi, Chuan-Yue Wang, Fei Wang, Kai Wang, Li Wang, Xiang Wang, Ying Wang, Xiao-Ping Wu, Xin-Ran Wu, Chun-Ming Xie, Guang-Rong Xie, Hai-Yan Xie, Peng Xie, Xiu-Feng Xu, Hong Yang, Jian Yang, Jia-Shu Yao, Shu-Qiao Yao, Ying-Ying Yin, Yong-Gui Yuan, Ai-Xia Zhang, Hong Zhang, Ke-Rang Zhang, Lei Zhang, Zhi-Jun Zhang, Ru-Bai Zhou, Yi-Ting Zhou, Jun-Juan Zhu, Chao-Jie Zou, Tian-Mei Si, Xi-Nian Zuo, Jing-Ping Zhao, Yu-Feng Zang

## Abstract

Major Depressive Disorder (MDD) is common and disabling, but its neuropathophysiology remains unclear. Most studies of functional brain networks in MDD have had limited statistical power and data analysis approaches have varied widely. The REST-meta-MDD Project of resting-state fMRI (R-fMRI) addresses these issues. Twenty-five research groups in China established the REST-meta-MDD Consortium by contributing R-fMRI data from 1,300 patients with MDD and 1,128 normal controls (NCs). Data were preprocessed locally with a standardized protocol prior to aggregated group analyses. We focused on functional connectivity (FC) within the default mode network (DMN), frequently reported to be increased in MDD. Instead, we found decreased DMN FC when we compared 848 patients with MDD to 794 NCs from 17 sites after data exclusion. We found FC reduction only in recurrent MDD, not in first-episode drug-naïve MDD. Decreased DMN FC was associated with medication usage but not with MDD duration. DMN FC was also positively related to symptom severity but only in recurrent MDD. Exploratory analyses also revealed alterations in FC of visual, sensory-motor and dorsal attention networks in MDD. We confirmed the key role of DMN in MDD but found reduced rather than increased FC within the DMN. Future studies should test whether decreased DMN FC mediates response to treatment. Finally, all resting-state fMRI indices of data contributed by the REST-meta-MDD consortium are being shared publicly via the R-fMRI Maps Project.

**SIGNIFICANCE STATEMENT:** Functional connectivity within the default mode network in major depressive disorder patients has been frequently reported abnormal but with contradicting directions in previous small sample size studies. In creating the REST-meta-MDD consortium containing neuroimaging data of 1,300 depressed patients and 1,128 normal controls from 25 research groups in China, we found decreased default mode network functional connectivity in depressed patients, driven by patients with recurrent depression, and associated with current medication treatment but not with disease duration. These findings suggest that default mode network functional connectivity remains a prime target for understanding the pathophysiology of depression, with particular relevance to revealing mechanisms of effective treatments.

## 1. INTRODUCTION

Major Depressive Disorder (MDD) is the second leading-cause of disability world-wide, with point prevalence exceeding 4% (1). The pathophysiology of MDD remains unknown despite intensive efforts, including neuroimaging studies. However, the small sample size of most MDD neuroimaging studies entails low sensitivity and reliability (2, 3). An exception is the Enhancing NeuroImaging Genetics through Meta-Analysis (ENIGMA) consortium which meta- and mega-analyzed thousands of structural MRI scans from MDD patients and healthy controls (4, 5). The ENIGMA-MDD working group found a slight albeit robust reduction in hippocampal volume (4) and cortical thinning in medial orbitofrontal cortex (5). However, this approach does not consider communication among brain regions, i.e., functional brain networks.

Abnormal communication among functional brain networks has been reported in MDD using resting-state fMRI (R-fMRI) functional connectivity (FC), which detects synchronized spontaneous activity among anatomically distinct networks. MDD studies have focused on the default mode network (DMN), which has been linked to rumination (6). The first study focusing on the DMN in MDD reported increased DMN FC (7), although similar studies found both increased and decreased DMN FC in MDD (8, 9). Meta-analyses have reported increased DMN FC in MDD, albeit based on few studies (6, 10). As summarized in SI Appendix, Table S1, 38 studies have examined DMN FC alterations in MDD. Of these, 18 found increases, eight decreases, seven both increases and decreases, and five no significant changes. As shown in SI Appendix, Figure S1, a voxel-wise meta-analysis of 32 studies revealed increased orbitofrontal DMN FC and decreased FC between dorsomedial prefrontal cortex (dmPFC) and posterior DMN in MDD. Such complex results may have contributed to prior inconsistencies.

Inconsistencies may reflect limited statistical power (2) from small samples, but data analysis flexibility may also contribute, as a large number of preprocessing and analysis operations with many different parameter combinations have been used in fMRI analyses (11). MDD studies have used diverse multiple comparison correction methods, most likely inadequate (12). Data analysis flexibility also impedes large-scale meta-analysis (6, 10). Moreover, clinical characteristics such as number and type of episodes, medication status and illness duration vary across studies, further contributing to heterogeneous results.

To address limited statistical power and analytic heterogeneity, we initiated the REST-meta-MDD Project. We implemented a standardized preprocessing protocol on Data Processing Assistant for Resting-State fMRI (DPARSF) (13) at local sites with only final indices provided to the consortium. We obtained R-fMRI indices (including FC matrices) corresponding to 1,300 patients with MDD and 1,128 normal controls (NCs) from 25 cohorts in China. To our knowledge, REST-meta-MDD is the largest MDD R-fMRI database (see SI Appendix, Table S2). We used linear mixed models to identify abnormal FC patterns associated with DMN across cohorts, and investigated whether episode type, medication status, illness severity and illness duration contributed to abnormalities.

## 2. RESULTS

### 2.1. Sample Composition

Contributions were requested from users of DPARSF, a MATLAB- and SPM-based R-fMRI preprocessing pipeline (13). Twenty-five research groups from 17 hospitals in China formed the REST-meta-MDD consortium and agreed to share final R-fMRI indices from patients with MDD and matched normal controls (see SI Appendix, Table S3 for data composition; henceforth “site” refers to each cohort for convenience) from studies approved by local Institutional Review Boards. The consortium contributed 2428 previously collected datasets (1300 MDDs and 1128 NCs) (Figure 1 and SI Appendix, Tables S3-5). On average, each site contributed 52.0±52.4 patients with MDD (range 13-282) and 45.1±46.9 NCs (range 6-251). Most MDD patients were female (826 vs. 474 males), as expected. The 562 patients with first episode MDD included 318 first episode drug-naïve (FEDN) MDD and 160 scanned while receiving antidepressants (medication status unavailable for 84). Of 282 with recurrent MDD, 121 were scanned while receiving antidepressants and 76 were not being treated with medication (medication status unavailable for 85). Episodicity (first or recurrent) and medication status were unavailable for 456 patients.

**Figure 1.**
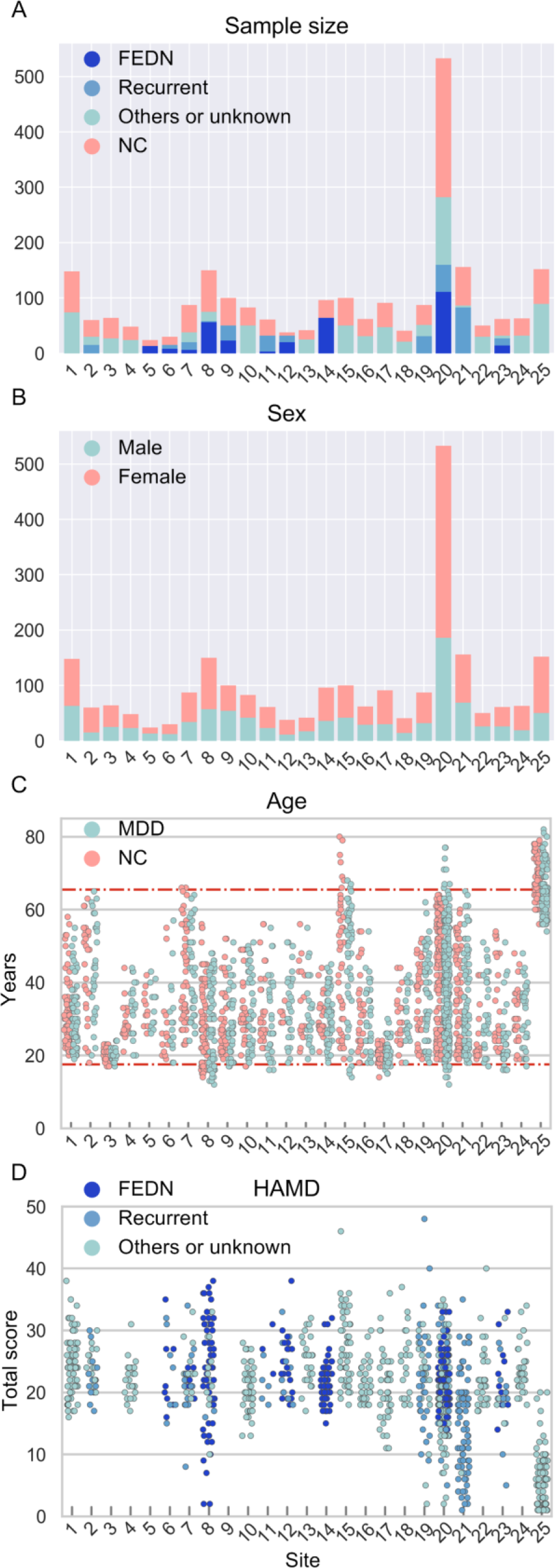
REST-meta-MDD sample characteristics. (A) Total number of participants per group for each contributing site. The MDD patients were subdivided into first-episode drug-naïve (FEDN), recurrent and others/unknown types. (B) Number of male subjects and female subjects for each site. (C) Age (in years) for all individuals per site for the MDD group and NC group. The two horizontal lines represents ages 18 and 65, the age limits for participants chosen for imaging analysis. (D) The score of Hamilton Depression Rating Scale (HAMD) for MDD patients, when available.

### 2.2. Decreased DMN FC in MDD Patients

Individual-level imaging processing was performed at each site using standardized DPARSF processing parameters. After preprocessing, time-series for the Dosenbach 160 functional regions-of-interest (ROIs) (14) were extracted. Individual-level imaging metrics (i.e., ROI time-series and R-fMRI indices) and phenotypic data were then uploaded through the R-fMRI Maps Project (http://rfmri.org/maps) platform at the Institute of Psychology, Chinese Academy of Sciences for statistical analyses. We defined DMN ROIs as those overlapping with the DMN delineated by Yeo et al (15). Average FC within the 33 DMN ROIs was taken to represent DMN within-network FC. We used the Linear Mixed Model (LMM) (16) to compare MDDs with NCs while allowing the effect to vary across sites. Mean DMN within-network FC (averaged across 33*32/2=528 connections) was compared between 848 MDDs and 794 NCs (see Sample Selection in SI Appendix, Supplementary Methods) with the LMM. MDD patients demonstrated significantly lower DMN within-network FC than NCs (T=−3.762, P=0.0002, d=−0.186, Figure 2A). On subgroup analyses, FEDN MDDs did not differ significantly from NCs (T=−0.914, P=0.361, d=−0.076, Figure 2B), while DMN FC was significantly decreased in patients with recurrent MDD vs. NCs (T=−3.737, P=0.0002, d=−0.326, Figure 2C). Significantly reduced DMN FC in recurrent MDD patients directly compared to FEDN MDDs (T=−2.676, P=0.008, d=−0.400, Figure 2D) suggests the recurrent MDDs were the major contributors to decreased DMN FC in MDD.

**Figure 2.**
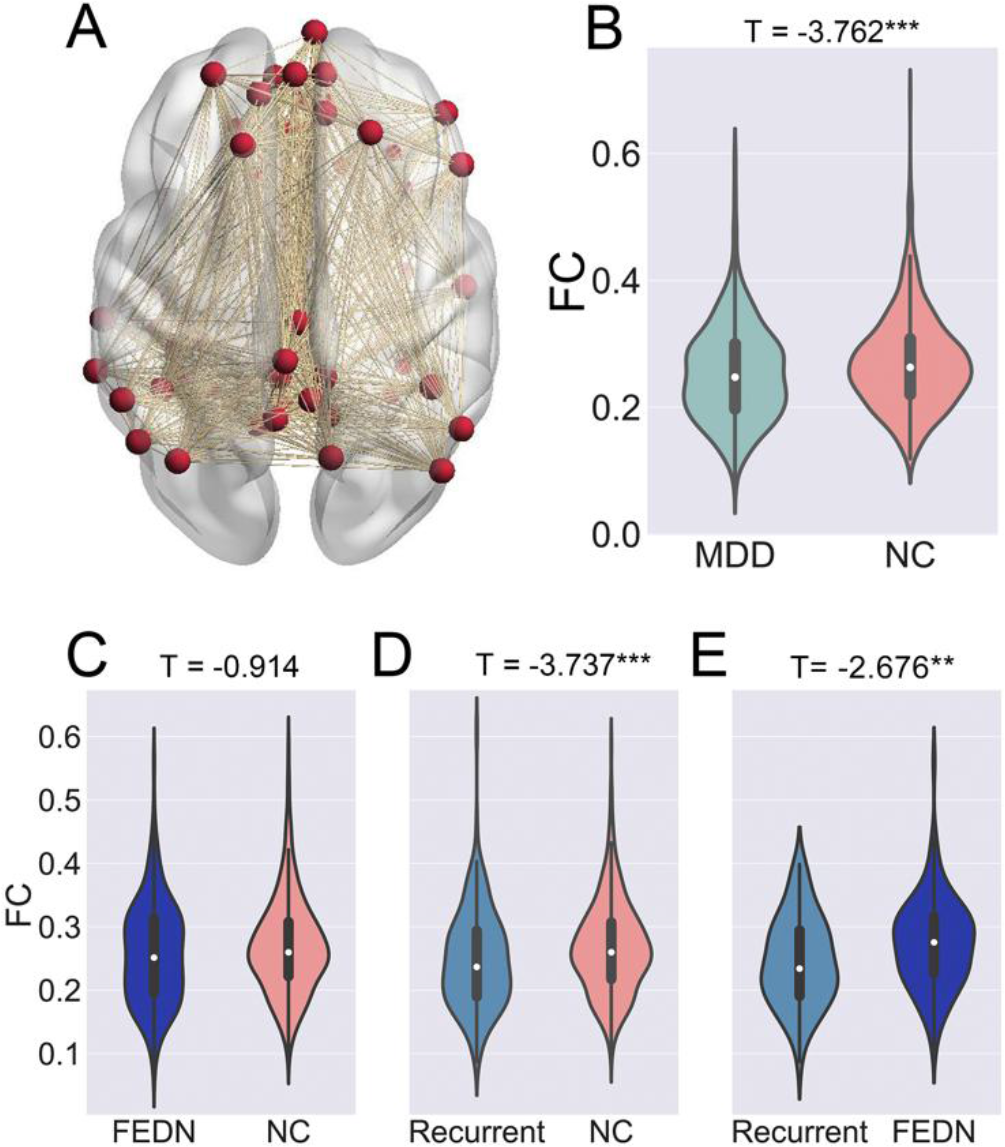
Decreased DMN functional connectivity in MDD patients. Mean DMN within-network FC was averaged across 33*32/2=528 connections as shown in A. The violin figures show the distribution of mean DMN within-network FC contrasting: MDD and NC groups (B); first episode drug naïve (FEDN) MDD and NC groups (C); recurrent MDD and NC groups (D); and FEDN MDD and recurrent MDD groups (E). Of note, for each comparison, only sites with sample size larger than 10 in each group were included. The T values were the statistics for these comparisons in Linear Mixed Model analyses. Please see Figure S3 for the forest plots of effect size per site generated by a meta-model in reproducibility analyses. **, p < 0.01; ***, p < 0.001.

### 2.3. Reduced DMN FC Was Not Associated with Illness Duration

Reduced DMN FC in recurrent MDD but not in FEDN MDD could reflect illness duration or medication history. We first tested the effect of illness duration in FEDN MDDs to reduce medication confounds. The tercile with longest illness duration (≥12 months, 70 MDDs from 2 sites) did not differ significantly from the tercile with shortest illness duration (≤3 months, 48 MDDs from the same 2 sites) in DMN FC (T=1.140, P=0.257, d=0.214, Figure 3A). Similarly, when exploring in the entire sample, the tercile with longest illness duration (≥24 months, 186 MDDs from 4 sites) did not differ significantly from the tercile with shortest illness duration (≤6 months, 112 MDDs from the same 4 sites): T=1.541, P=0.124, d=0.184 (Figure 3B). Beyond chronicity, clinical subtypes could contribute to DMN FC. We examined subtypes characterized by core depression, anxiety, and neurovegetative symptoms of melancholia by mapping HAMD scale items to National Institute of Mental Health Research-Domain-Criteria (RDoC) constructs (17). However, subtype analyses did not reveal any significant effects (see SI Appendix, Supplementary Results, Table S6, Figures S5 and S6).

**Figure 3.**
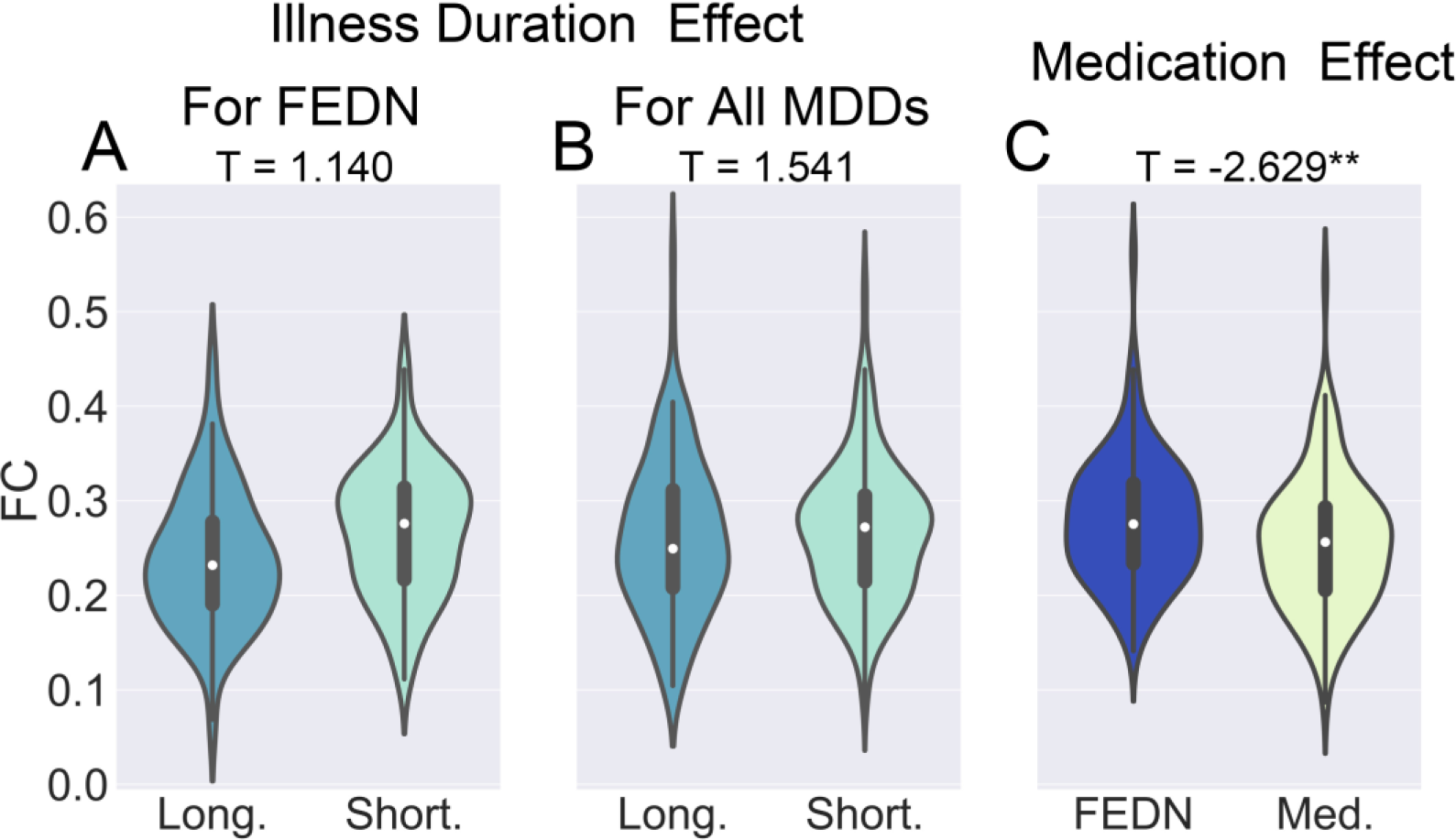
The effects of illness duration and medication status on decreased DMN functional connectivity in MDD patients. The violin figures show the distribution of mean DMN within-network FC for first episode drug naïve (FEDN) MDD with long vs. short illness duration (A), for all MDD patients with long vs. short illness duration (B), and for first episode MDD patients with vs. without medication usage (C). The T values are the statistics for these comparisons in Linear Mixed Model analyses. Please see Figure S4 for the forest plots of effect size per site generated by a meta-model in reproducibility analyses. **, p < 0.01.

### 2.4. Medication Effect and Reduced DMN FC in MDD Patients

To further examine medication treatment effects, we contrasted first episode MDDs on medication (115 MDDs from Site 20) with FEDN MDDs (97 MDDs from Site 20) and found significantly reduced DMN FC (T=−2.629, P=0.009, d=−0.362, Figure 3C). When directly comparing 102 first episode MDDs on medication with 266 NCs from 2 sites, we found a non-significant effect (T=−1.614, P=0.108, d=−0.188). While FEDN MDDs showed higher DMN FC than recurrent MDDs as shown in Section 2.2, 102 first-episode MDDs on medication and 57 recurrent MDDs from 2 sites did not differ significantly (T=0.548, P=0.585, d=−0.091). This suggests that medication treatment might account for our overall finding of reduced DMN FC in MDD. However, we could not address whether currently unmedicated recurrent MDDs had been previously treated with antidepressants. We were also unable to examine treatment duration, as medication status was binary.

### 2.5. Association of DMN FC with Symptom Severity

The association between DMN FC and HAMD scores was tested on 734 MDD patients (excluding remitted patients with HAMD scores below 7) from 15 sites and was not significant (T=1.591, P=0.112, r=0.059). The effect of symptom severity was not significant in FEDN MDDs (N=197, 3 sites; T=−0.158, P=0.874, r=−0.011), but significant in recurrent MDDs (N=126, 4 sites; T=2.167, P=0.032, r=0.194).

### 2.6. Reproducibility

We assessed reproducibility through several strategies (SI Appendix, Table S7). 1) Using another functional clustering atlas generated by parcellating whole brain R-fMRI data into spatially coherent regions of homogeneous FC (i.e., Craddock’s 200 functional clustering atlas (18), with 48 DMN ROIs) confirmed our results, except that the effect of symptom severity in recurrent MDDs became insignificant (T=1.424, P=0.157, r=0.129). 2) Using a finer-grade parcellations (i.e., Zalesky’s random 980 parcellation (19), with 211 DMN ROIs) also confirmed our results, except that symptom severity in recurrent MDDs became insignificant (T=1.264, P=0.209, r=0.115). 2) Beyond LMM, we also performed meta-analyses: within-site T-values were converted into Hedge’s *g*, and entered in a random effect meta-model (using R “metansue”, https://www.metansue.com/). Results were almost the same, although the difference between recurrent MDDs and FEDN MDDs became insignificant (Z=−1.732, P=0.083, d=−0.251), and symptom severity in recurrent MDDs became insignificant (Z=1.304, P=0.192, r=0.119). 3) We also tested whether global signal regression (GSR) mattered. With GSR, we found similar results except for loss of significance for the difference between recurrent MDDs and FEDN MDDs (T=−0.974, P=0.331, d=−0.145), the medication effect (T=−1.891, P=0.060, d=−0.261), and symptom severity in recurrent MDD (T=1.741, P=0.084, r=0.157). This overall confirmation is important since the global signal has been viewed as reflecting spurious noise (20), and its standard deviation differed significantly between MDDs and NCs (T=−2.662, P=0.008, d=−0.131). 4) For head motion control, despite already incorporating the Friston-24 model at the individual level and a motion covariate at the group level in primary analyses, we also used scrubbing (removing time points with framewise displacement >0.2mm (21) to verify results. All results remained the same using this aggressive head motion control strategy.

### 2.7. Exploratory Findings of Brain Networks Beyond DMN

Although we focused on DMN FC in MDD, we also performed exploratory analyses comprising other brain networks beyond DMN using the 7-network atlas developed by Yeo et al. (15): visual network (VN), sensory-motor network (SMN), dorsal attention network (DAN), ventral attention network (VAN), subcortical network (instead of the limbic network defined by Yeo et al., which is not covered by the 160 ROIs), frontoparietal network (FPN) and DMN. Comparing all 848 MDDs with 794 NCs, after false discovery rate (FDR) correction among 7 within-network and 21 between-network connections, we found VN, SMN, and DMN demonstrated decreased within-network connection in MDDs as compared to NC. Furthermore, 3 between-network connections also demonstrated significant decreases in MDDs: VN-SMN, VN-DAN, and SMN-DAN (Figure 4A, SI Appendix, Table S8). We further explored which subgroups contributed to these 6 abnormal within- and between-network connections by performing subgroup analyses. FEDN MDDs only demonstrated significant decrease in within-network connectivity of VN after FDR correction (Figure 4B). Recurrent MDDs demonstrated the same abnormal pattern as the whole group, confirming again they were the major contributors (Figure 4C). This was further supported by the direct comparisons between recurrent MDDs with FEDN MDDs, which showed lower within-network connectivity of DMN and between-network connectivity of VN-SMN and SMN-DAN in recurrent MDDs (Figure 4D, SI Appendix, Table S9). Similar to the primary DMN analysis, we did not find any significant illness duration effect, whether within the whole group or within FEDN MDDS (SI Appendix, Table S9). When comparing MDDs on medication with FEDN MDDs, reduced within-network connectivity of DMN and between-network connectivity of SMN and DAN was found in MDDs with medication (Figure 4E). Finally, none of the within- and between-network connectivities correlated significantly with illness severity (HAMD) after correction (SI Appendix, Table S10).

**Figure 4.**
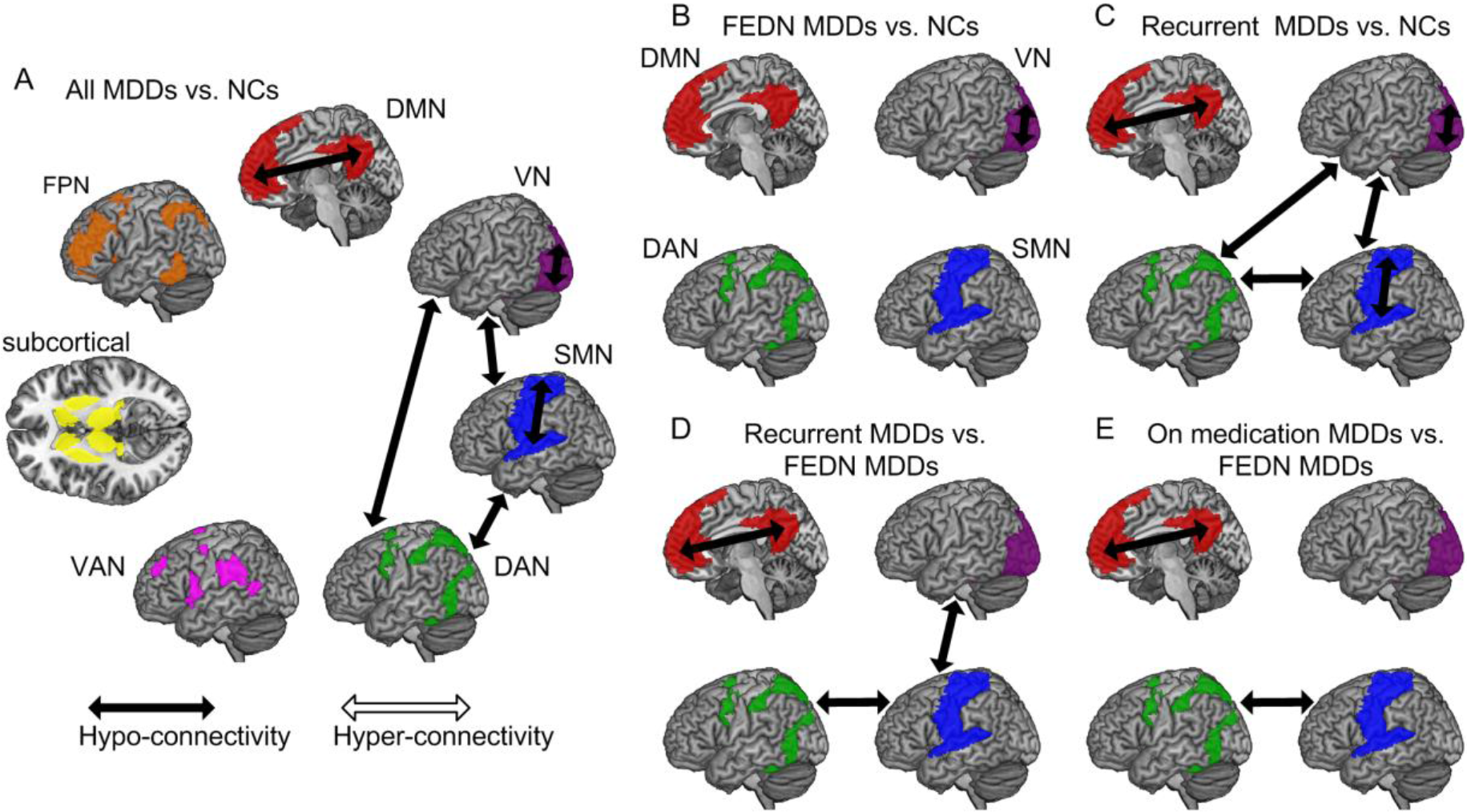
Exploratory analyses of functional connectivity within- and between-the 7 brain networks delineated by Yeo et al. (15): (A) All MDDs vs. NCs; (B) FEDN MDDs vs. NCs; (C) Recurrent MDDs vs. NCs; (D) Recurrent MDDs vs. FEDN MDDs; (E) MDDs on medication vs. FEDN MDDs. False discovery rate (FDR) correction was performed among 7 within-network and 21 between-network connections for the whole-group analysis (comparing all 848 MDDs with 794 NCs). For subgroup analyses, FDR corrected for the 6 abnormal connections found in the whole-group analysis. VN, visual network; SMN: sensory-motor network; DAN: dorsal attention network; VAN: ventral attention network; Subcortical: subcortical ROIs; FPN: frontal parietal network; DMN: default mode network.

## 3. DISCUSSION

Using an unprecedentedly large sample, we found decreased instead of increased FC within the DMN in MDD compared with NCs. However, this effect was only significant in recurrent MDD whether vs. controls or patients with FEDN MDD. Furthermore, decreased DMN FC in recurrent MDD was associated with being scanned on antidepressant medication rather than illness duration. DMN FC was also positively related to symptom severity but only in recurrent MDD. Exploratory analyses revealed increased ReHo in left DLPFC in FEDN MDD, and decreased ReHo in bilateral primary motor cortex in recurrent MDD.

Our primary results contradict the prevailing notion that DMN FC is increased in MDD (6, 10). Several factors may account for this discrepancy. 1) Prior studies have also reported decreased DMN FC in MDD (see SI Appendix, Table S1). Our voxel-wise meta-analysis of 32 studies (SI Appendix, Figure S1) revealed both increases (orbitofrontal DMN FC) and decreases (dmPFC / posterior DMN FC) in MDD. 2) Prior inconsistent results may also reflect heterogeneous analysis strategies (11). We applied a standardized analysis protocol across sites, removing analytic variations. 3) Average DMN FC might be insensitive to possible pair-wise increases in MDD DMN FC. However, pair-wise tests did not reveal even a single pair of significantly increased within-DMN connection in MDDs, even within the three DMN subsystems proposed by Andrews-Hanna et al. (22) (see SI Appendix, Supplementary Results and Figure S7). Finally, most studies reporting increased DMN FC in MDDs, albeit inconsistently, were conducted in Caucasian samples, while our sample was homogeneously Chinese. Ethnic differences may have contributed, as east Asians report lower lifetime prevalence of MDD (1), more somatic symptoms and fewer psychological symptoms (23), and differ in MDD risk genes (24). International studies will need to address this question.

In subgroup analyses, we only found decreased DMN FC in recurrent MDD patients, with nearly twice the effect size of the whole-group (d=−0.326 vs. −0.186). Similarly, ENIGMA-MDD found a robust reduction in hippocampal volume (a key DMN node) only in recurrent MDD and not in first episode MDD (4). Illness duration in recurrent MDD was significantly longer than in FEDN MDD (Z=6.419, p<0.001), but it was unrelated to DMN FC on direct comparisons. An early MDD study (7) found that DMN FC was positively correlated with current episode duration but this was not confirmed subsequently (9, 25). We conclude that illness duration is likely unrelated to DMN FC. However, longitudinal studies are needed to determine whether DMN FC changes over the course of depressive episodes.

Decreased DMN FC in recurrent MDD was associated with antidepressant medication treatment. We confirmed that first episode MDDs scanned while on medication had decreased DMN FC than FEDN MDD. This result aligns with studies of antidepressants on DMN FC in MDD (26), dysthymia (27), and in healthy individuals (28). In MDD, antidepressant treatment for 12 weeks reduced posterior DMN FC (26). In patients with dysthymia, 10 weeks of duloxetine treatment reduced DMN FC (27). In healthy individuals, duloxetine for 2 weeks reduced DMN FC and improved mood (28). Our finding of medication-associated reduction in DMN FC suggests antidepressant medications may alleviate depressive symptoms by reducing DMN FC. This medication effect (effect size d=−0.362) might also underlie the contradiction between our finding of reduced DMN FC in MDD and prior meta-analyses. However, this medication effect was observed in a retrospective cross-sectional sample that cannot be stratified by class, dosage, or length of use, thus has it to be confirmed using longitudinal designs with medication follow-up.

We did not find significant associations between DMN FC and symptom severity in all MDDs nor in FEDN MDDs. However, symptom severity was positively correlated with DMN FC in recurrent MDDs. Similarly, a prior report (29) found a positive correlation between DMN FC in a specific frontal subcircuit and illness severity in MDDs (half treated with medication). Our finding may reflect medication effects in recurrent MDD (the effect was stronger in recurrent MDDs on medication: N=40, 2 sites; T=3.268, P=0.003, r=0.489): the greater the medication benefit (indicated by lower HAMD score), the more DMN FC was reduced. However, this finding should be interpreted with caution, as these small sample size secondary analyses might not reflect a true effect (2). Additionally, this result was not consistently confirmed with other parcellations (see SI Appendix, Table S7). More importantly, testing this hypothesis requires longitudinal follow-up of medication effects.

To extend beyond the DMN, we explored other brain networks defined by Yeo et al. (15). We found decreased FC within VN, SMN and DMN. Task-based fMRI studies have reported abnormal neural filtering of irrelevant visual information in visual cortex in MDD (30). R-fMRI studies have also found reduced VN FC in MDD patients (31), suggesting abnormal processing in the visual cortex in MDD. For SMN, a previous meta-analysis (32) reported reduced regional homogeneity in depressed patients, which could underlie psychomotor retardation, a core clinical manifestation of MDD (33). Besides changes in within-network FC, we also observed decreased between-network FC involving VN, SMN and DAN. The reduced FC of the SMN with the VN and DMN may be interpreted as the neural underpinnings of the pervasive influence of psychomotor retardation on attentional processes, as revealed by previous studies (34). Similar to the primary analyses focused on the DMN, most of these other alterations in FC were contributed by recurrent MDD patients, which needs to be confirmed by future longitudinal designs.

Study limitations include an exclusively Chinese sample, with unknown generalization to other populations. As a next step, we plan to analyze UK Biobank MDD data (35). In addition, in conjunction with the ENIGMA-MDD consortium (36), we are inviting international MDD researchers to join the REST-meta-MDD Project to identify ethnicity/culture-general and ethnicity/culture-specific abnormal brain patterns in MDD. Second, we could not address longitudinal effects, such as response to treatment. We anticipate the REST-meta-MDD consortium will perform coordinated prospective longitudinal studies. Third, medication treatment was binary; future studies should quantify cumulative doses and include non-pharmacologic treatments. Finally, our findings require independent replication (11). To improve transparency and reproducibility, the analysis code has been openly shared at https://github.com/Chaogan-Yan/PaperScripts/tree/master/Yan_2018. Upon publication, the R-fMRI indices of the 1300 MDD patients and 1128 NCs will be openly shared through the R-fMRI Maps Project (LINK_TO_BE_ADDED). These data derivatives will allow replication, secondary analyses and discovery efforts while protecting participant privacy and confidentiality. Future independent efforts could include generating neural biotypes of MDD (37), performing dynamic FC analysis and data mining with machine learning algorithms.

In summary, based on the largest R-fMRI database of MDD, we confirmed the key role of the DMN in MDD, identifying a reduction of DMN FC in patients with recurrent MDD. This reduction appears to reflect medication usage rather than illness duration. These findings suggest that the DMN should remain a prime target for further MDD research, especially to determine whether reducing DMN FC mediates symptomatic improvement.

## 4. MATERIALS AND METHODS

### 4.1. Phenotypic Data

Consortium members (25 research groups from 17 Chinese hospitals) met on March 25^th^, 2017 to establish the collaboration; all agreed to provide diagnosis, age at scan, sex and education. When collected systematically, measures of first-episode or recurrent MDD (if a patient’s prior and current episode were diagnosed as MDD based on ICD10 or DSM-IV), medication status, illness duration, 17-item Hamilton Depression Rating Scale (HAMD) were also provided.

### 4.2. Individual-Level Image Processing

Neuroimaging analysts from each site took a two-day DPARSF training course on May 13-14, 2017 at the Institute of Psychology, Chinese Academy of Sciences to harmonize analyses of individual R-fMRI data and 3D T1-weighted images.

#### 4.2.1. DMN FC Analyses

After preprocessing (SI Appendix, Supplementary Methods), time-series for the Dosenbach 160 functional regions-of-interest (ROIs) (14) were extracted. Dosenbach 160 functional ROIs were used for the primary analysis as these functionally defined regions were based on a series of five meta-analyses, focused on error-processing, default-mode (task-induced deactivations), memory, language and sensorimotor functions. For each, we defined DMN ROIs as those overlapping with the DMN delineated by Yeo et al. (15) The average FC (Fisher’s r-to-z transformed Pearson’s correlation between time-series of all ROI pairs) within DMN ROIs was defined as DMN within-network FC for patient-control contrasts.

### 4.3. Group-Level Image Processing

#### 4.3.1. Sample Selection

From 1300 MDDs and 1128 NCs, we selected 848 MDDs and 794 NCs from 17 sites for statistical analyses. Exclusion criteria (e.g., incomplete information, bad spatial normalization, bad coverage, excessive head motion and sites with fewer than 10 subjects in either group) and final inclusions are provided in SI Appendix, Supplementary Methods and Figure S2.

#### 4.3.2. Statistical Analyses

We used the Linear Mixed Model (LMM) to compare MDDs with NCs while allowing site-varying effects. LMM describes the relationship between a response variable (e.g., DMN FC) and independent variables (here diagnosis and covariates of age, sex, education, and head motion), with coefficients that can vary with respect to grouping variables (here site) (16). We utilized MATLAB’s command fitlme (https://www.mathworks.com/help/stats/fitlme.html) to test the model: y ~ 1 + Diagnosis + Age + Sex + Education + Motion + (1 | Site) + (Diagnosis | Site), which yields T and P values for the fixed effect of Diagnosis. Cohen’s d effect size was computed as 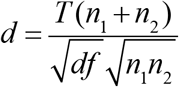 (38).

#### 4.3.3. Subgroup Analyses

Several sites reported whether patients with MDD were in their first episode (and drug-naïve) or recurrent. We compared 232 FEDN MDD patients with 394 corresponding NCs from 5 sites. We also compared 189 recurrent MDD patients with 427 corresponding NCs from 6 sites. To compare 119 FEDN MDD patients with 72 recurrent MDD patients from 2 sites, we replaced Diagnosis with FEDN or recurrent status in the LMM model.

#### 4.3.4. Analyses of Effects of Illness Duration, Medication, and Symptom Severity

As the distribution of illness duration was skewed (most were brief), we contrasted the terciles with longest and shortest illness durations instead of Diagnosis in the LMM model. To test medication effects, we replaced Diagnosis with medication (on/off, assessed at time of scan) in the LMM model. Finally, to test symptom severity effects, we replaced Diagnosis with the 17-item HAMD total score regressor in the LMM model.

## ACKNOWLEDGEMENTS

This work was supported by the National Key R&D Program of China (2017YFC1309902), the National Natural Science Foundation of China (81671774, 81630031, 81471740 and 81371488), the Hundred Talents Program and the 13th Five-year Informatization Plan (XXH13505) of Chinese Academy of Sciences, Beijing Municipal Science & Technology Commission (Z161100000216152, Z171100000117016, Z161100002616023 and Z171100000117012), Department of Science and Technology, Zhejiang Province (2015C03037) and the National Basic Research (973) Program (2015CB351702).

## CONFLICTS OF INTEREST

All the authors declare no competing financial interests.

## REFERENCES

1. Ferrari AJ, et al. (2013) Burden of Depressive Disorders by Country, Sex, Age, and Year: Findings from the Global Burden of Disease Study 2010. PLOS Medicine 10(11):e1001547.

2. Button KS, et al. (2013) Power failure: why small sample size undermines the reliability of neuroscience. Nat Rev Neurosci 14(5):365–376.

3. Chen X, Lu B, & Yan CG (2018) Reproducibility of R-fMRI metrics on the impact of different strategies for multiple comparison correction and sample sizes. Hum Brain Mapp 39(1):300–318.

4. Schmaal L, et al. (2016) Subcortical brain alterations in major depressive disorder: findings from the ENIGMA Major Depressive Disorder working group. Mol Psychiatry 21(6):806–812.

5. Schmaal L, et al. (2017) Cortical abnormalities in adults and adolescents with major depression based on brain scans from 20 cohorts worldwide in the ENIGMA Major Depressive Disorder Working Group. Molecular psychiatry 22(6):900–909.

6. Hamilton JP, Farmer M, Fogelman P, & Gotlib IH (2015) Depressive Rumination, the Default-Mode Network, and the Dark Matter of Clinical Neuroscience. Biological psychiatry 78(4):224–230.

7. Greicius MD, et al. (2007) Resting-state functional connectivity in major depression: abnormally increased contributions from subgenual cingulate cortex and thalamus. Biological psychiatry 62(5):429–437.

8. Zhu X, et al. (2012) Evidence of a dissociation pattern in resting-state default mode network connectivity in first-episode, treatment-naive major depression patients. Biological psychiatry 71(7):611–617.

9. Guo W, et al. (2014) Abnormal default-mode network homogeneity in first-episode, drug-naive major depressive disorder. PLoS ONE 9(3):e91102.

10. Kaiser RH, Andrews-Hanna JR, Wager TD, & Pizzagalli DA (2015) Large-Scale Network Dysfunction in Major Depressive Disorder: A Meta-analysis of Resting-State Functional Connectivity. JAMA Psychiatry 72(6):603–611.

11. Poldrack RA, et al. (2017) Scanning the horizon: towards transparent and reproducible neuroimaging research. Nat Rev Neurosci 18(2):115–126.

12. Eklund A, Nichols TE, & Knutsson H (2016) Cluster failure: Why fMRI inferences for spatial extent have inflated false-positive rates. Proc Natl Acad Sci U S A.

13. Yan CG & Zang YF (2010) DPARSF: A MATLAB Toolbox for “Pipeline” Data Analysis of Resting-State fMRI. Front Syst Neurosci 4:13.

14. Dosenbach NU, et al. (2010) Prediction of individual brain maturity using fMRI. Science (New York, N.Y 329(5997):1358–1361.

15. Yeo BT, et al. (2011) The organization of the human cerebral cortex estimated by intrinsic functional connectivity. Journal of neurophysiology 106(3):1125–1165.

16. West BT, Welch KB, & Galecki AT (2014) Linear mixed models: a practical guide using statistical software (CRC Press).

17. Ahmed AT, et al. (2018) Mapping depression rating scale phenotypes onto research domain criteria (RDoC) to inform biological research in mood disorders. Journal of affective disorders 238:1–7.

18. Craddock RC, James GA, Holtzheimer PE, 3rd, Hu XP, & Mayberg HS (2012) A whole brain fMRI atlas generated via spatially constrained spectral clustering. Hum Brain Mapp 33(8):1914–1928.

19. Zalesky A, et al. (2010) Whole-brain anatomical networks: does the choice of nodes matter? Neuroimage 50(3):970–983.

20. Power JD, Plitt M, Laumann TO, & Martin A (2016) Sources and implications of whole-brain fMRI signals in humans. Neuroimage.

21. Jenkinson M, Bannister P, Brady M, & Smith S (2002) Improved Optimization for the Robust and Accurate Linear Registration and Motion Correction of Brain Images. NeuroImage 17(2):825–841.

22. Andrews-Hanna JR, Reidler JS, Sepulcre J, Poulin R, & Buckner RL (2010) Functional-anatomic fractionation of the brain’s default network. Neuron 65(4):550–562.

23. Ryder AG, et al. (2008) The cultural shaping of depression: somatic symptoms in China, psychological symptoms in North America? Journal of abnormal psychology 117(2):300–313.

24. Long H, et al. (2013) The long rather than the short allele of 5-HTTLPR predisposes Han Chinese to anxiety and reduced connectivity between prefrontal cortex and amygdala. Neurosci Bull 29(1):4–15.

25. Wise T, et al. (2017) Instability of default mode network connectivity in major depression: a two-sample confirmation study. Translational psychiatry 7(4):e1105.

26. Li B, et al. (2013) A treatment-resistant default mode subnetwork in major depression. Biological psychiatry 74(1):48–54.

27. Posner J, et al. (2013) Antidepressants normalize the default mode network in patients with dysthymia. JAMA Psychiatry 70(4):373–382.

28. van Wingen GA, et al. (2014) Short-term antidepressant administration reduces default mode and task-positive network connectivity in healthy individuals during rest. Neuroimage 88:47–53.

29. Davey CG, Harrison BJ, Yucel M, & Allen NB (2012) Regionally specific alterations in functional connectivity of the anterior cingulate cortex in major depressive disorder. Psychological medicine 42(10):2071–2081.

30. Desseilles M, et al. (2009) Abnormal neural filtering of irrelevant visual information in depression. J Neurosci 29(5):1395–1403.

31. Veer IM, et al. (2010) Whole brain resting-state analysis reveals decreased functional connectivity in major depression. Front Syst Neurosci 4.

32. Iwabuchi SJ, et al. (2015) Localized connectivity in depression: A meta-analysis of resting state functional imaging studies. Neuroscience & Biobehavioral Reviews 51:77–86.

33. Buyukdura JS, McClintock SM, & Croarkin PE (2011) Psychomotor retardation in depression: biological underpinnings, measurement, and treatment. Progress in neuro-psychopharmacology & biological psychiatry 35(2):395–409.

34. Snyder HR (2013) Major depressive disorder is associated with broad impairments on neuropsychological measures of executive function: a meta-analysis and review. Psychological bulletin 139(1):81–132.

35. Sudlow C, et al. (2015) UK biobank: an open access resource for identifying the causes of a wide range of complex diseases of middle and old age. PLoS medicine 12(3):e1001779.

36. Thompson PM, et al. (2014) The ENIGMA Consortium: large-scale collaborative analyses of neuroimaging and genetic data. Brain imaging and behavior 8(2):153–182.

37. Drysdale AT, et al. (2017) Resting-state connectivity biomarkers define neurophysiological subtypes of depression. Nat Med 23(1):28–38.

38. Rosenthal R & Rosnow RL (1991) Essentials of behavioral research: Methods and data analysis (3rd ed.) (McGraw-Hill, New York, NY).

