## Supplementary Materials for "Reduced Default Mode Network Functional Connectivity in Patients with Recurrent Major Depressive Disorder"

### SUPPLEMENTARY METHODS

**Privacy Protection**

Data submitted to the consortium were fully deidentified and anonymized. We utilized the following strategies to ensure the privacy of the subjects during data uploading: 1) None of the HIPAA identifiable elements were shared. The only shared information is: subject ID, sex, age, education, episode status, medication status, illness duration, Hamilton Depression Rating Scale (HAMD) and Hamilton Anxiety Rating Scale (HAMA). 2) The subject ID uploaded was reprogramed, so that it could not be traced back to the original subject ID used in the original studies. 3) There was no face information in the MRI data, as no original T1 image was shared.

**Preprocessing**

First, the initial 10 volumes were discarded, and slice-timing correction was performed. Then, the time series of images for each subject were realigned using a six-parameter (rigid body) linear transformation. After realignment, individual T1-weighted images were co-registered to the mean functional image using a 6 degrees-of-freedom linear transformation without re-sampling and then segmented into gray matter (GM), white matter (WM) and cerebrospinal fluid (CSF) (1). Finally, transformations from individual native space to MNI space were computed with the Diffeomorphic Anatomical Registration Through Exponentiated Lie algebra (DARTEL) tool (2).

**Nuisance Regression**

To minimize head motion confounds, we utilized the Friston 24-parameter model (3) to regress out head motion effects. Additionally, mean framewise displacement (FD, derived from Jenkinson's relative root mean square algorithm) (4) was used to address the residual effects of motion as a covariate in group analyses. In validation analysis, scrubbing (removing time points with FD > 0.2mm) was also utilized to verify results using an aggressive head motion control strategy. As global signal regression (GSR) is still a controversial practice in the R-fMRI field (5), we did not perform GSR in primary analyses, but included analyses with GSR for validation. Other sources of spurious variance (WM and CSF signals) were also removed from the data through linear regression to reduce respiratory and cardiac effects. Additionally, linear trend were included as a regressor to account for drifts in the blood oxygen level dependent (BOLD) signal. We performed temporal bandpass filtering (0.01-0.1Hz) on all time series.

**Exploratory Analyses of a Broad Array of R-fMRI Metrics**

Beyond the hypothesis-driven analysis of default mode network (DMN) functional connectivity (FC), we also shared whole-brain voxel-wise R-fMRI metrics for exploring local abnormalities of major depressive disorder (MDD).

Amplitude of Low Frequency Fluctuations (ALFF) (6) and fractional ALFF (fALFF) (7): ALFF is the mean of amplitudes within a specific frequency domain (here, 0.01-0.1Hz) from a fast Fourier transform of a voxel's time course. fALFF is a normalized version of ALFF and represents the relative contribution of specific frequency band oscillations to the whole detectable frequency range. Of note, preprocessing of temporal bandpass filtering (0.01-0.1Hz) was not performed for ALFF/fALFF analyses.

Regional Homogeneity (ReHo) (8): ReHo is a rank-based Kendall's coefficient of concordance (KCC) that assesses the synchronization among time courses of nearest neighboring voxels (here, 27 voxels).

Degree Centrality (DC) (9, 10): DC is the number or sum of weights of significant connections for a voxel. Here, we calculated the weighted sum of positive correlations by requiring each connection's correlation coefficient to exceed a threshold of $r > 0.25$ (9).

Voxel-mirrored homotopic connectivity (VMHC) (11, 12): VMHC corresponds to the functional connectivity between any pair of symmetric inter-hemispheric voxels - that is, the Pearson's correlation coefficient between the time series of each voxel and that of its counterpart voxel at the same location in the opposite hemisphere. The resultant VMHC values were Fisher-Z transformed. For better correspondence between symmetric voxels, VMHC requires that individual functional data to be further registered to a symmetric template and smoothed (4 mm FWHM). The group averaged symmetric template was created by first computing a mean normalized T1 image across participants, and then this image was averaged with its left–right mirrored version (12).

Before entering into further analyses, all of the metric maps were Z-standardized (subtracting the mean value of the entire brain from each voxel, and dividing by the corresponding standard deviation) and then smoothed (4 mm FWHM), except for VMHC (which were smoothed and Fisher's r-to-z transformed beforehand).

**Sample Selection**

From 1300 MDDs and 1128 NCs, we selected subjects for group statistical analyses through the following criteria (please also see Supplementary Figure S2): 1) Site 25 was excluded since it mainly contained late onset depression (most with age > 60) and remitted patients, resulting in 1211 MDDs and 1064 NCs; 2) subjects without information on sex, age and education were excluded, resulting in 1150 MDDs and 971 NCs; 3) subjects with bad imaging data and bad spatial normalization (by visual inspection) were excluded, resulting in 1042 MDDs and 884 NCs; 4) subjects with age less than 18 or more than 65 were excluded, resulting in 989 MDDs and 860 NCs; 5) subjects with bad coverage (<90% of the group mask) or excessive head motion (mean FD > 0.2mm) were excluded, resulting in 943 MDDs and 846 NCs; 6) to further remove subjects with distortions that not screened by visual inspection, we excluded subjects with spatial correlation < 0.6 (a threshold defined by mean - 2SD) between each participant's ReHo map and the group mean ReHo map, resulting in 900 MDDs and 815 NCs; 7) finally, we removed those sites with fewer than 10 subjects in either group (10 was selected arbitrarily to balance the objectives of optimizing overall sample size and minimizing extreme biases), resulting in 848 MDDs and 794 NCs from 17 sites.

#### SUPPLEMENTARY RESULTS

##### Effects of Demographic Covariates

In the LMM model of statistical analyses, we have included several demographic covariates to control their confounding effects (i.e., age, sex, education, and head motion). All these covariates showed significant effects on DMN FC, thus confirmed the necessity for controlling them. Females demonstrated stronger DMN FC than males (T = 2.130, p = 0.033, d = 0.018), which is consistent with many prior studies (13, 14). DMN FC decreased as age increased (T = -6.297, $p < 10^{-9}$, r = -0.154), and also decreased when more education was achieved (T = -3.338, p = 0.0009, r = -0.082). Head motion also has a strong impact on DMN FC (T = 10.513, $p < 10^{-24}$, r = -0.252). We have also tested interactions between Diagnosis and these demographic covariates (by entering the interaction term into the LMM model once upon a time), none of them was significant (Diagnosis*Age: T = 0.112, p = 0.911; Diagnosis*Sex: T = 0.339, p = 0.734; Diagnosis*Education: T = -0.239, p = 0.811; Diagnosis*Head Motion: T = -0.696, p = 0.487).

##### Effects of Clinical Subtypes

Recently, Ahmed et al. defined three clinical subtypes by mapping the HAMD scale to the National Institute of Mental Health Research-Domain-Criteria (RDoC) constructs: Core Depression (CD), Anxiety (ANX), and Neurovegetative Symptoms of Melancholia (NVSM) (15). Here we investigated the impact of these three subtypes on DMN FC (we could not define the fourth subtype of Atypical Depression since we lacked data on the Quick Inventory of Depressive Symptomatology, similar to

Ahmed et al.'s Emory PReDICT sample). No significant difference in DMN FC was found between CD+ and CD- (T = 0.431, p = 0.667), ANX+ and ANX- (T = 0.477, p = 0.634), as well as between NVSM+ and NVSM- (T = -0.029, p = 0.977) (Supplementary Table S6 and Figure S5). We further performed pairwise contrasts of DMN FC among CD+, ANX+ and NVSM+ subgroups, while excluding those comorbid for the subtypes being examined. However, we did not find any significant differences in these subtype comparisons: CD+ vs. ANX+ (T = -0.941, p = 0.349), CD+ vs. NVSM+ (T = 1.457, p = 0.147) and ANX+ vs. NVSM+ (T = 1.072, p = 0.285) (Supplementary Table S6 and Figure S6).

**Connection-wise Analysis of DMN FC in MDD**

In the primary analysis, we averaged FC across the 528 (i.e., 33*32/2) pairs of 33 DMN ROIs (Dosenbach's template) to represent the overall DMN FC. This overall measure might be over-simplified and insensitive to changes of single connections. Thus, in supplementary analyses, we also compared pair-wise connection within these DMN ROIs to identify which pair of DMN ROIs contributed the most. False discovery rate (FDR) multiple comparison correction strategy was utilized to correct the comparisons of 528 pairs. For the comparison of FC within the DMN between MDDs with NCs, 42 pairs of within-DMN connections showed significantly decreased FC in the MDDs, while none displayed increased FC (Supplementary Figure S7A and Supplementary Table S11). The decreased FC mainly involved the regions of ventromedial prefrontal cortex (vmPFC), posterior cingulate cortex (PCC), superior frontal gyrus, lateral temporal cortex (LTC), and inferior parietal lobe (IPL) bilaterally. Interestingly, across the 528 connections within the DMN, the abnormality extent of DMN connections (quantified by T-value of difference between 848 MDDs and 794 NCs) was negatively correlated with the strength of those connections (quantified by averaging FC strength across the subjects): r = -0.513, p < 0.001. That means the stronger the DMN connection, the higher probability that it was affected (decreased) in MDDs. We further divided the MDDs into first episode drug naïve (FEDN) MDDs and recurrent MDDs, as we did in the main text. For the comparison of FC between recurrent MDDs with NCs, 24 pairs of within-DMN connections showed significantly decreased FC in the recurrent MDDs, while none displayed the reverse effect (Supplementary Figure S7B and Supplementary Table S12). The decreased FC mainly involved the regions of vmPFC, PCC, LTC, and IPL bilaterally, which largely overlapped with those identified in the analysis including all MDDs. In contrast, for the comparison of FC between FEDN MDDs with NCs, none of the within-DMN connections showed significant difference of FC (i.e., survived FDR correction, Supplementary Figure S7C and Supplementary Table S12). In addition, the direct comparison of FC between FEDN MDDs with recurrent MDDs revealed 3 pairs of connection with lower FC in the recurrent MDDs, which involved the dorsomedial PFC, ACC, bilateral angular gyrus, and left LTC (Supplementary Figure S7D and Supplementary Table S12). These results from group comparisons of connection-by-connection FC were in accord with those reported in the main text for which the FC was averaged across the DMN. To further confirm these results, we tested the FCs grouped into 3 DMN subsystems as proposed by Andrews-Hanna et al (16, 17): 1) core subsystem, 2) dorsal medial prefrontal cortex (dmPFC) subsystem, and 3) medial temporal lobe (MTL) subsystem. ROIs overlapped with the corresponding Yeo's 17 networks (18) were assigned to the subsystem as dissected by Andrews-Hanna et al. (2014) (17). As demonstrated in Supplementary Table S13, none of the subsystem demonstrated the effects in the reversed direction as the main results, while most effects were focused in the core and dmPFC subsystems. Together, all of these results indicated that recurrent MDD patients, but not FEDN MDD patients, demonstrated decreased DMN FC, as compared to NCs.

We also tested the effects of illness duration and medication on pair-wise connection results. The comparison of DMN FC between FEDN MDDs with longer illness duration and those with shorter illness duration revealed no significant difference on any connection after FDR correction. The effect of illness duration was also not significant for all MDD patients. In contrast, the comparison of DMN FC between first episode MDDs on medication with FEDN MDDs showed that medication usage was associated with decreased FC between left PCC and right PCC (Supplementary Figure S7E). These results were consistent with the overall DMN FC analysis reported in the main text.

**Local Abnormalities in MDD**

Although we focused on FC in the primary analysis, we also performed exploratory analyses to illustrate the potential value of the shared voxel-wise R-fMRI metric maps to reveal local abnormalities. Since regional homogeneity (ReHo) demonstrated the highest test-retest reliability among commonly used R-fMRI indices (19), we present ReHo abnormalities in MDD and include other indices as well: amplitude of low frequency fluctuations (ALFF), fractional ALFF (fALFF), degree centrality (DC), and voxel-mirrored homotopic connectivity (VMHC). In applying LMM voxel-wise, we used Gaussian random field theory to correct for multiple comparisons, with strict two-tailed thresholds (voxel $p<0.0005$ [$Z>3.29$] and cluster $p<0.025$), maintaining the family-wise error rate under 5% (20, 21). Comparing all 848 MDDs with 794 NCs, ReHo was increased in left dorsolateral prefrontal cortex (DLPFC) in MDD and decreased in bilateral primary motor cortex (Supplementary Figure S8A). Among subgroups, left DLPFC ReHo was significantly increased in FEDN MDDs (Supplementary

Figure S8B) but not in recurrent MDD (Supplementary Figure S8C). By contrast, ReHo in bilateral primary motor cortex was only significantly decreased in recurrent MDDs (Supplementary Figure S8C) but not in FEDN MDDs (Supplementary Figure S8B). However, FEDN MDDs and recurrent MDDs did not differ significantly in ReHo when compared directly.

As depicted in Supplementary Figures S9-S12, significantly lower ALFF, fALFF, DC and VMHC in MDDs compared to NCs were found in precuneus or PCC. These areas are believed to be key nodes of DMN (22, 23), and have been found to be altered in MDD patients (24). Further analysis revealed that this effect might be largely driven by the difference between recurrent MDDs and NCs (Supplementary Figures S10 and S11). Besides, IPL, another DMN key region, was also found to be significantly altered in MDD (Supplementary Figures S9, S10, and S12). Interestingly, we found FEDN MDDs' VMHC and DC were significantly higher than recurrent MDDs in the middle temporal lobe, which is also an important DMN region. Given these two regional metrics' similarities to FC, these results further supported our main findings: MDDs are characterized by lower FCs within DMN, and this effect is largely driven by recurrent MDDs.

**REFERENCES**


1. Ashburner J & Friston KJ (2005) Unified segmentation. *NeuroImage* 26(3):839-851.
2. Ashburner J (2007) A fast diffeomorphic image registration algorithm. *NeuroImage* 38(1):95-113.
3. Friston KJ, Williams S, Howard R, Frackowiak RSJ, & Turner R (1996) Movement-Related effects in fMRI time-series. *Magn Reson Med* 35(3):346-355.
4. Jenkinson M, Bannister P, Brady M, & Smith S (2002) Improved Optimization for the Robust and Accurate Linear Registration and Motion Correction of Brain Images. *NeuroImage* 17(2):825-841.
5. Murphy K & Fox MD (2016) Towards a Consensus Regarding Global Signal Regression for Resting State Functional Connectivity MRI. *Neuroimage*.
6. Zang YF*, et al.* (2007) Altered baseline brain activity in children with ADHD revealed by resting-state functional MRI. *Brain Dev* 29(2):83-91.
7. Zou Q-H*, et al.* (2008) An improved approach to detection of amplitude of low-frequency fluctuation (ALFF) for resting-state fMRI: Fractional ALFF. *J Neurosci Methods* 172(1):137-141.
8. Zang Y, Jiang T, Lu Y, He Y, & Tian L (2004) Regional homogeneity approach to fMRI data

analysis. *NeuroImage* 22(1):394-400.

9. Buckner RL, *et al.* (2009) Cortical hubs revealed by intrinsic functional connectivity: mapping, assessment of stability, and relation to Alzheimer's disease. *J Neurosci* 29(6):1860-1873.

10. Zuo XN, *et al.* (2012) Network Centrality in the Human Functional Connectome. *Cereb Cortex* 22(8):1862-1875.

11. Anderson JS, *et al.* (2011) Decreased interhemispheric functional connectivity in autism. *Cerebral cortex* 21(5):1134-1146.

12. Zuo XN, *et al.* (2010) Growing together and growing apart: regional and sex differences in the lifespan developmental trajectories of functional homotopy. *J Neurosci* 30(45):15034-15043.

13. Biswal BB, *et al.* (2010) Toward discovery science of human brain function. *Proc Natl Acad Sci U S A* 107(10):4734-4739.

14. Yan CG, Craddock RC, Zuo XN, Zang YF, & Milham MP (2013) Standardizing the intrinsic brain: towards robust measurement of inter-individual variation in 1000 functional connectomes. *Neuroimage* 80:246-262.

15. Ahmed AT, *et al.* (2018) Mapping depression rating scale phenotypes onto research domain criteria (RDoC) to inform biological research in mood disorders. *J Affect Disord* 238:1-7.

16. Andrews-Hanna JR, Reidler JS, Sepulcre J, Poulin R, & Buckner RL (2010) Functional-anatomic fractionation of the brain's default network. *Neuron* 65(4):550-562.

17. Andrews-Hanna JR, Smallwood J, & Spreng RN (2014) The default network and self-generated thought: component processes, dynamic control, and clinical relevance. *Ann N Y Acad Sci* 1316(1):29-52.

18. Yeo BT, *et al.* (2011) The organization of the human cerebral cortex estimated by intrinsic functional connectivity. *J Neurophysiol* 106(3):1125-1165.

19. Zuo XN & Xing XX (2014) Test-retest reliabilities of resting-state FMRI measurements in human brain functional connectomics: a systems neuroscience perspective. *Neurosci Biobehav Rev* 45:100-118.

20. Eklund A, Nichols TE, & Knutsson H (2016) Cluster failure: Why fMRI inferences for spatial extent have inflated false-positive rates. *Proc Natl Acad Sci U S A*.

21. Chen X, Lu B, & Yan CG (2018) Reproducibility of R-fMRI metrics on the impact of different strategies for multiple comparison correction and sample sizes. *Hum Brain Mapp* 39(1):300-318.

22. Raichle ME, *et al.* (2001) A default mode of brain function. *Proceedings of the National Academy of Sciences* 98(2):676-682.

23. Raichle ME & Snyder AZ (2007) A default mode of brain function: a brief history of an evolving idea. *Neuroimage* 37(4):1083-1090; discussion 1097-1089.

24. Zhu X, *et al.* (2012) Evidence of a dissociation pattern in resting-state default mode network connectivity in first-episode, treatment-naive major depression patients. *Biol Psychiatry* 71(7):611-617.

25. Greicius MD, *et al.* (2007) Resting-state functional connectivity in major depression: abnormally increased contributions from subgenual cingulate cortex and thalamus. *Biological psychiatry* 62(5):429-437.

26. Bluhm R, *et al.* (2009) Resting state default-mode network connectivity in early depression using a seed region-of-interest analysis: Decreased connectivity with caudate nucleus. *Psychiatry and Clinical Neurosciences* 63(6):754-761.

27. Cullen KR, *et al.* (2009) A preliminary study of functional connectivity in comorbid adolescent depression. *Neuroscience Letters* 460(3):227-231.

28. Sheline YI, Price JL, Yan Z, & Mintun MA (2010) Resting-state functional MRI in depression unmasks increased connectivity between networks via the dorsal nexus. *Proc Natl Acad Sci U S A* 107(24):11020-11025.

29. Zhou Y, *et al.* (2010) Increased neural resources recruitment in the intrinsic organization in major depression. *J Affect Disord* 121(3):220-230.

30. Berman MG, *et al.* (2011) Depression, rumination and the default network. *Social cognitive and affective neuroscience* 6(5):548-555.

31. Wu M, *et al.* (2011) Default-mode network connectivity and white matter burden in late-life depression. *Psychiatry Res* 194(1):39-46.

32. Alexopoulos GS, *et al.* (2012) Functional connectivity in the cognitive control network and the default mode network in late-life depression. *J Affect Disord* 139(1):56-65.

33. Davey CG, Harrison BJ, Yucel M, & Allen NB (2012) Regionally specific alterations in functional connectivity of the anterior cingulate cortex in major depressive disorder. *Psychol Med* 42(10):2071-2081.

34. Lui S, *et al.* (2011) Resting-state functional connectivity in treatment-resistant depression. *Am J Psychiatry* 168(6):642-648.

35. Peng DH, *et al.* (2012) Abnormal functional connectivity with mood regulating circuit in unmedicated individual with major depression: a resting-state functional magnetic resonance study. *Chinese medical journal* 125(20):3701-3706.

36. Andreescu C, *et al.* (2013) Resting state functional connectivity and treatment response in

late-life depression.

37. de Kwaasteniet B, *et al.* (2013) Relation between structural and functional connectivity in major depressive disorder. *Biol Psychiatry* 74(1):40-47.

38. Li B, *et al.* (2013) A treatment-resistant default mode subnetwork in major depression. *Biol Psychiatry* 74(1):48-54.

39. Guo W, *et al.* (2014) Abnormal default-mode network homogeneity in first-episode, drug-naive major depressive disorder. *PLoS One* 9(3):e91102.

40. Connolly CG, *et al.* (2013) Resting-state functional connectivity of subgenual anterior cingulate cortex in depressed adolescents. *Biol Psychiatry* 74(12):898-907.

41. Liston C, *et al.* (2014) Default mode network mechanisms of transcranial magnetic stimulation in depression. *Biol Psychiatry* 76(7):517-526.

42. Manoliu A, *et al.* (2013) Insular dysfunction within the salience network is associated with severity of symptoms and aberrant inter-network connectivity in major depressive disorder. *Frontiers in human neuroscience* 7:930.

43. Sambataro F, Wolf ND, Pennuto M, Vasic N, & Wolf RC (2014) Revisiting default mode network function in major depression: evidence for disrupted subsystem connectivity. *Psychol Med* 44(10):2041-2051.

44. van Tol MJ, *et al.* (2014) Local cortical thinning links to resting-state disconnectivity in major depressive disorder. *Psychol Med* 44(10):2053-2065.

45. Crowther A, *et al.* (2015) Resting-state connectivity predictors of response to psychotherapy in major depressive disorder. *Neuropsychopharmacology* 40(7):1659-1673.

46. Chen Y, Wang C, Zhu X, Tan Y, & Zhong Y (2015) Aberrant connectivity within the default mode network in first-episode, treatment-naive major depressive disorder. *J Affect Disord* 183:49-56.

47. Sawaya H, *et al.* (2015) Resting-state functional connectivity of antero-medial prefrontal cortex sub-regions in major depression and relationship to emotional intelligence. *The international journal of neuropsychopharmacology* 18(6).

48. Eyre HA, *et al.* (2016) Altered resting-state functional connectivity in late-life depression: A cross-sectional study. *J Affect Disord* 189:126-133.

49. Goya-Maldonado R, *et al.* (2016) Differentiating unipolar and bipolar depression by alterations in large-scale brain networks. *Hum Brain Mapp* 37(2):808-818.

50. Kim SM, *et al.* (2016) Affective network and default mode network in depressive adolescents with disruptive behaviors. *Neuropsychiatr Dis Treat* 12:49-56.

51. Deng D, *et al.* (2016) Modulation of the Default Mode Network in First-Episode, Drug-Naive Major Depressive Disorder via Acupuncture at Baihui (GV20) Acupoint. *Frontiers in human neuroscience* 10:230.

52. Parlar M, *et al.* (2017) Relation between patterns of intrinsic network connectivity, cognitive functioning, and symptom presentation in trauma-exposed patients with major depressive disorder. *Brain and behavior* 7(5):e00664.

53. Straub J, *et al.* (2017) Successful group psychotherapy of depression in adolescents alters fronto-limbic resting-state connectivity. *J Affect Disord* 209:135-139.

54. Zhu X, Zhu Q, Shen H, Liao W, & Yuan F (2017) Rumination and Default Mode Network Subsystems Connectivity in First-episode, Drug-Naive Young Patients with Major Depressive Disorder. *Sci Rep* 7:43105.

55. Schilbach L, *et al.* (2016) Transdiagnostic commonalities and differences in resting state functional connectivity of the default mode network in schizophrenia and major depression. *Neuroimage Clin* 10:326-335.

56. Yang R, *et al.* (2016) Decreased functional connectivity to posterior cingulate cortex in major depressive disorder. *Psychiatry research. Neuroimaging* 255:15-23.

57. Wang Y, *et al.* (2017) Topologically convergent and divergent functional connectivity patterns in unmedicated unipolar depression and bipolar disorder. *Transl Psychiatry* 7(7):e1165.

58. Evans JW, *et al.* (2018) Default Mode Connectivity in Major Depressive Disorder Measured Up to 10 Days After Ketamine Administration. *Biol Psychiatry*.

59. Knyazev GG, *et al.* (2018) Task-positive and task-negative networks in major depressive disorder: A combined fMRI and EEG study. *J Affect Disord* 235:211-219.

60. Xia M, *et al.* (2019) Reproducibility of functional brain alterations in major depressive disorder: Evidence from a multisite resting-state functional MRI study with 1,434 individuals. *Neuroimage* 189:700-714.

61. Penninx BW, *et al.* (2008) The Netherlands Study of Depression and Anxiety (NESDA): rationale, objectives and methods. *International journal of methods in psychiatric research* 17(3):121-140.

62. Pannekoek JN, *et al.* (2015) Investigating distinct and common abnormalities of resting-state functional connectivity in depression, anxiety, and their comorbid states. *European neuropsychopharmacology : the journal of the European College of Neuropsychopharmacology* 25(11):1933-1942.

63. Teismann H, *et al.* (2014) Establishing the bidirectional relationship between depression and

subclinical arteriosclerosis--rationale, design, and characteristics of the BiDirect Study. *BMC psychiatry* 14:174.

64. Teuber A, *et al.* (2017) MR imaging of the brain in large cohort studies: feasibility report of the population- and patient-based BiDirect study. *European radiology* 27(1):231-238.
65. Sundermann B, *et al.* (2017) Diagnostic classification of unipolar depression based on resting-state functional connectivity MRI: effects of generalization to a diverse sample. *Journal of neural transmission (Vienna, Austria : 1996)* 124(5):589-605.
66. Feder S, *et al.* (2017) Sample heterogeneity in unipolar depression as assessed by functional connectivity analyses is dominated by general disease effects. *Journal of Affective Disorders* 222:79-87.
67. Vogelbacher C, *et al.* (2018) The Marburg-Munster Affective Disorders Cohort Study (MACS): A quality assurance protocol for MR neuroimaging data. *Neuroimage* 172:450-460.
68. Redlich R, *et al.* (2014) Brain morphometric biomarkers distinguishing unipolar and bipolar depression. A voxel-based morphometry-pattern classification approach. *JAMA psychiatry* 71(11):1222-1230.
69. Yuksel D, *et al.* (2018) Neural correlates of working memory in first episode and recurrent depression: An fMRI study. *Progress in neuro-psychopharmacology & biological psychiatry* 84(Pt A):39-49.
70. Sudlow C, *et al.* (2015) UK Biobank: An Open Access Resource for Identifying the Causes of a Wide Range of Complex Diseases of Middle and Old Age. *PLOS Medicine* 12(3):e1001779.
71. Shen X, *et al.* (2017) Subcortical volume and white matter integrity abnormalities in major depressive disorder: findings from UK Biobank imaging data. *Sci Rep* 7(1):5547.
72. Wigmore EM, *et al.* (2017) Do regional brain volumes and major depressive disorder share genetic architecture? A study of Generation Scotland (n=19 762), UK Biobank (n=24 048) and the English Longitudinal Study of Ageing (n=5766). *Transl Psychiatry* 7(8):e1205.
73. Consortium GNC (2014) The German National Cohort: aims, study design and organization. *European Journal of Epidemiology* 29(5):371-382.
74. Teipel SJ, *et al.* (2017) Multicenter stability of resting state fMRI in the detection of Alzheimer's disease and amnestic MCI. *Neuroimage Clin* 14:183-194.
75. Yahata N, *et al.* (2016) A small number of abnormal brain connections predicts adult autism spectrum disorder. *Nature communications* 7:11254.
76. Ichikawa N, *et al.* (2017) Identifying melancholic depression biomarker using whole-brain functional connectivity.

77. Schmaal L*, et al.* (2016) Cortical abnormalities in adults and adolescents with major depression based on brain scans from 20 cohorts worldwide in the ENIGMA Major Depressive Disorder Working Group. *Mol Psychiatry*.

78. Schmaal L*, et al.* (2016) Subcortical brain alterations in major depressive disorder: findings from the ENIGMA Major Depressive Disorder working group. *Mol Psychiatry* 21(6):806-812.

79. Wang L*, et al.* (2013) Interhemispheric functional connectivity and its relationships with clinical characteristics in major depressive disorder: a resting state fMRI study. *PLoS One* 8(3):e60191.

80. Wang L*, et al.* (2015) The effects of antidepressant treatment on resting-state functional brain networks in patients with major depressive disorder. *Hum Brain Mapp* 36(2):768-778.

81. Liu Y*, et al.* (2017) Regional homogeneity associated with overgeneral autobiographical memory of first-episode treatment-naive patients with major depressive disorder in the orbitofrontal cortex: A resting-state fMRI study. *J Affect Disord* 209:163-168.

82. Guo W*, et al.* (2017) Decreased interhemispheric coordination in the posterior default-mode network and visual regions as trait alterations in first-episode, drug-naive major depressive disorder. *Brain imaging and behavior*.

83. Peng D*, et al.* (2014) Altered brain network modules induce helplessness in major depressive disorder. *Journal of Affective Disorders* 168:21-29.

84. Peng D*, et al.* (2015) Dissociated large-scale functional connectivity networks of the precuneus in medication-naïve first-episode depression. *Psychiatry Research: Neuroimaging* 232(3):250-256.

85. Zhu J*, et al.* (2014) Default-mode network connectivity in depression: A resting-state fMRI study (in Chinese). *Chinese Journal of Nervous and Mental Diseases* 40(8):454-458.

86. Shen Y*, et al.* (2015) Sub-hubs of baseline functional brain networks are related to early improvement following two-week pharmacological therapy for major depressive disorder. *Hum Brain Mapp* 36(8):2915-2927.

87. Tang Y*, et al.* (2013) Decreased functional connectivity between the amygdala and the left ventral prefrontal cortex in treatment-naive patients with major depressive disorder: a resting-state functional magnetic resonance imaging study. *Psychological Medicine* 43(9):1921-1927.

88. Li HJ*, et al.* (2014) Surface-based regional homogeneity in first-episode, drug-naive major depression: a resting-state FMRI study. *Biomed Res Int* 2014:374828.

89. Du L*, et al.* (2016) Changes in Problem-Solving Capacity and Association With Spontaneous

Brain Activity After a Single Electroconvulsive Treatment in Major Depressive Disorder. *The journal of ECT* 32(1):49-54.

90. Wu X, *et al.* (2016) Dysfunction of the cingulo-opercular network in first-episode medication-naive patients with major depressive disorder. *J Affect Disord* 200:275-283.

91. Yang X-h, *et al.* (2017) Anhedonia correlates with abnormal functional connectivity of the superior temporal gyrus and the caudate nucleus in patients with first-episode drug-naive major depressive disorder. *Journal of Affective Disorders* 218:284-290.

92. Hou Z, *et al.* (2018) Increased temporal variability of striatum region facilitating the early antidepressant response in patients with major depressive disorder. *Progress in neuro-psychopharmacology & biological psychiatry* 85:39-45.

93. Hou Z, *et al.* (2018) Distinctive pretreatment features of bilateral nucleus accumbens networks predict early response to antidepressants in major depressive disorder. *Brain imaging and behavior* 12(4):1042-1052.

94. Chen T, *et al.* (2017) Anomalous single-subject based morphological cortical networks in drug-naive, first-episode major depressive disorder. *Hum Brain Mapp* 38(5):2482-2494.

95. Cao J, *et al.* (2016) Resting-state functional MRI of abnormal baseline brain activity in young depressed patients with and without suicidal behavior. *J Affect Disord* 205:252-263.

96. Wang J, *et al.* (2017) Electroconvulsive therapy selectively enhanced feedforward connectivity from fusiform face area to amygdala in major depressive disorder. *Social cognitive and affective neuroscience* 12(12):1983-1992.

97. Cheng W, *et al.* (2016) Medial reward and lateral non-reward orbitofrontal cortex circuits change in opposite directions in depression. *Brain* 139(Pt 12):3296-3309.

98. Ye M, *et al.* (2015) Changes of Functional Brain Networks in Major Depressive Disorder: A Graph Theoretical Analysis of Resting-State fMRI. *PLOS ONE* 10(9):e0133775.

99. Luo Q, *et al.* (2015) Frequency Dependant Topological Alterations of Intrinsic Functional Connectome in Major Depressive Disorder. *Scientific Reports* 5:9710.

100. Xue S, Wang X, Wang W, Liu J, & Qiu J (2016) Frequency-dependent alterations in regional homogeneity in major depression. *Behavioural Brain Research* 306:13-19.

101. Zheng H, *et al.* (2018) The dynamic characteristics of the anterior cingulate cortex in resting-state fMRI of patients with depression. *Journal of Affective Disorders* 227:391-397.

102. Jing B, *et al.* (2013) Difference in amplitude of low-frequency fluctuation between currently depressed and remitted females with major depressive disorder. *Brain Research* 1540:74-83.

103. Yang X, *et al.* (2015) Anatomical and functional brain abnormalities in unmedicated major

depressive disorder. *Neuropsychiatr Dis Treat* 11:2415-2423.

104. Cheng Y, *et al.* (2017) Resting-state brain alteration after a single dose of SSRI administration predicts 8-week remission of patients with major depressive disorder. *Psychological Medicine* 47(3):438-450.

105. Yuan Y, *et al.* (2008) Abnormal neural activity in the patients with remitted geriatric depression: A resting-state functional magnetic resonance imaging study. *Journal of Affective Disorders* 111(2):145-152.

106. Radua J, *et al.* (2012) A new meta-analytic method for neuroimaging studies that combines reported peak coordinates and statistical parametric maps. *European psychiatry : the journal of the Association of European Psychiatrists* 27(8):605-611.

**SUPPLEMENTARY TABLES**

**Supplementary Table S1.** A summary of fMRI studies revealing altered default mode network (DMN) functional connectivity (FC) in individuals with MDD.

| Study | Sample Size | | MDD Group's Age, Mean (SD) | Methodology | Principle Findings on FC within DMN | | Multiple Comparison Correction Strategy | Episodes† |
|---|---|---|---|---|---|---|---|---|
| | MDD | Healthy Control | | | Increased FC | Decreased FC | | |
| Greicius et al., 2007* | 28 | 20 | 38.5 (N/A) | ICA | sgACC | | Joint expected probability distribution with height and extent thresholds of $p < 0.01$, masked within the DMN | N/A |
| Bluhm et al., 2009* | 14 | 15 | 21.9 (5.1) | Seed-based Analysis: PCC | no results | no results | FDR | N/A |
| Cullen et al., 2009 | 12 | 14 | 16.5 (0.95) | Seed-based Analysis: sgACC | | sgACC, right medial frontal cortex | GRF theory base correction (min $z > 2.3$, cluster significance: $p < 0.05$) | N/A |
| Sheline et al., 2010 | 18 | 17 | 35.9 (1.3) | Seed-based Analysis: PCC | dmPFC | | Thresholded using $p < 0.01$ (z = 2.58) | Eight first-episode drug naïve |
| Zhou et al., 2010* | 20 | 18 | 40.6 (10.7) | Seed-based Analysis: PCC | sgACC | | Within combined FC mask, $p < 0.01$ for each voxel and a cluster size of at least 675 $mm^3$, equal to the corrected threshold of $p < 0.001$, determined by a Monte Carlo simulation (AFNI AlphaSim) | All first-episode drug naïve |

| | | | | | | | | |
|---|---|---|---|---|---|---|---|---|
| Berman et al., 2011* | 15 | 15 | 25.7 (N/A) | Seed-based Analysis: PCC | sgACC | | $p < 0.001$ at the voxel-level and a cluster-size threshold of 26 voxels to produce a $p < 0.05$ threshold | N/A |
| Lui et al., 2011* | 32 | 48 | 32.0 (10) | Seed-based Analysis: ACC | | Left middle temporal gyrus, right inferior frontal gyrus | GRF theory base correction (p < 0.05, more than 5 contiguous voxels) | N/A |
| Wu et al., 2011 | 12 | 12 | 70.5 (4.9) | Seed-based Analysis: PCC | dmPFC | sgACC | Monte Carlo simulations (AFNI AlphaSim) using a small volume correction | Five first-episode drug naïve |
| Alexopoulos et al., 2012* | 16 | 10 | 67.9 (4.7) | Seed-based Analysis: PCC | vmPFC, dmPFC, medial temporal regions | | GRF theory base correction （$p < 0.01$ voxel wised and $p < 0.05$ cluster wised | N/A |
| Davey et al., 2012* | 18 | 20 | 18.9 (2.3) | Seed-based Analysis: sgACC | mPFC | | thresholded at $p < 0.001$, cluster-wise corrected ($p_{FWE} < 0.05$) | Nine first-episode drug naïve |
| Peng et al., 2012* | 16 | 16 | 33.4 (5.8) | Seed-based Analysis: pgACC | parahippocampus gyrus | | $p < 0.05$ voxel-wise | N/A |

| | | | | | | | | |
|---|---|---|---|---|---|---|---|---|
| Zhu et al., 2012* | 35 | 35 | 20.5 (1.8) | ICA | dmPFC, pgACC | PCC/PCu, bilateral AG | $p < 0.05$ with FDR correction, masked within the DMN | All first-episode drug naïve |
| Andreescu et al., 2013* | 47 | 46 | 68.7 (7.0) | Seed-based Analysis: PCC | PCu | | Small volume multiple correction embedded in SPM5 (voxel $p < 0.001$) | Six first-episode drug naïve |
| Connolly et al., 2013* | 30 | 45 | 16 (0.3) | Seed-based Analysis: sgACC | bilateral inferior frontal gyrus, right parahippocampal, bilateral Inferior parietal lobule, right superior temporal gyrus | | A Monte Carlo simulation with a voxel-wise threshold, p <0.05 | N/A |
| De Kwaasteniet et al., 2013* | 18 | 24 | 44.6 (10.4) | Seed-based Analysis: sgACC | MTL (uncorrected) | | FWE correction ($p < .05$) | N/A |
| Li et al., 2013 | 24 | 24 | 31.8 (11.1) | ICA | mPFC, PCC | | $p < 0.05$, family-wise error corrected | 1.83 |
| Guo et al., 2014* | 24 | 24 | 25.6 (7.5) | ICA | dmPFC | rLTC | $p < 0.05$ for multiple comparisons according to GRF theory (min $z > 1.96$, cluster significance: $p < 0.05$) | All first-episode drug naïve |
| Liston et al., 2014* | 17 | 35 | 42.3 (17.3) | Seed-based Analysis: sgACC | mPFC, PCu | | A cluster threshold (K > 16 voxels, $p < 0.01$ for network-of-interest analyses; K > 25, $p < 0.005$ for whole brain analyses) | N/A |
| Manoliu et al., 2014* | 25 | 25 | 48.8 (14.9) | ICA | mPFC, PCu | PCu | $p < 0.05$, FWE-corrected | 5.56 |

| | | | | | | | | |
|---|---|---|---|---|---|---|---|---|
| Sambataro et al., 2014* | 20 | 20 | 33.6 (11.0) | ICA | PCC, sgACC, retrosplenial PCC, right lateral temporo-parietal cortex | | FDR ($q < 0.05$) | N/A |
| van Tol et al., 2014* | 20 | 20 | 38.3 (11.6) | Seed-based Analysis: dmPFC | | PCC, AG, LTG | correction level $p < 0.05$ FWE-corrected, initial voxel-wise threshold $p = 0.001$ | Five first-episode drug naïve |
| Chen et al., 2015* | 38 | 38 | 32.1 (7.7) | Seed-based Analysis: PCC | | dmPFC, right inferior parietal gyrus/AG | FDR ($q < 0.05$) | All first-episode drug naïve |
| Crowther et al., 2015* | 23 | 20 | 33.09 (7.45) | Seed-based Analysis: MPFC, PCC | Left middle temporal lobe | | Voxel wise Z > 2.3, Cluster wise p < 0.05 | 3.39 (1.8) |
| Pannekoek et al., 2015* | 37 | 48 | 35.7 (10.11) | ICA with dual regression | no results | no results | PT with TFCE | N/A |
| Peng et al., 2015* | 16 | 16 | 34.43 (6.72) | Seed-based Analysis: PCC | | Right middle temporal gyrus, Right superior temporal gyrus, Left inferior parietal gyrus, Left superior temporal gyrus, Right superior medial frontal gyrus, | FWER corrected: voxel wise p < 0.05, cluster wise p < 0.05 | All first-episode drug naïve |

| | | | | | | | | |
|---|---|---|---|---|---|---|---|---|
| | | | | | | Right anterior cingulate gyrus, Left middle frontal gyrus | | |
| Sawaya et al., 2015* | 21 | 21 | 37.3 (14.2) | Seed-based Analysis: sgACC | | ACC, mPFC | GRF (*p* < 0.01 voxel wised, *p* < 0.05 cluster wised) | N/A |
| Deng et al., 2016* | 29 | 29 | 28.69 (6.69) | Seed-based Analysis: PCC | Middle prefrontal cortex, Angular gyrus | ACC | FDR | FEDN |
| Eyre et al., 2016 | 17 | 31 | 67.3 (6.6) | ICA | pSTS | | A single-voxel threshold of $z > 1.64$, $p < 0.05$, with correction for cluster extent using Random Field Theory at $p < 0.05$ | N/A |
| Goya-Maldonado et al., 2016* | 20 | 20 | 35.6 (10.4) | ICA | parahippocampus gyrus, PCC/PCu | | Voxel threshold $p < 0.005$, cluster size $k > 48$ (AlphaSim implemented in REST) | 4.2 |
| Kim et al., 2016* | 22 | 20 | 13.9 (1.6) | Seed-based Analysis: PCC | inferior parietal lobe | | FWE corrected. An uncorrected $p < 0.001$ and 40 extended voxels | All first-episode drug naïve |
| Schilbach et al., 2016* | 102 | 106 | 37.75 (13.29) | Seed-based Analysis: MPFC, PCC | | PCC: right superior parietal cortex, left superior parietal cortex; MPFC: left precentral gyrus | voxel wise p < 0.001 and cluster wise p < 0.05 | NAN |
| Yang et al., 2016 | 23 | 25 | 33.4 (9.7) | Seed-based | | temporal lobe | A threshold of p < 0.001 and extended clusters of 4 20 voxels | N/A |

| | | | | | | | | |
|---|---|---|---|---|---|---|---|---|
| | | | | Analysis: PCC | | | | |
| Parlar et al., 2017* | 21 | 20 | 40.2 (17.9) | ICA | mPFC | | FDR-corrected | 13.2 |
| Straub et al., 2017* | 19 | 19 | 16.6 (1.4) | Seed-based Analysis: sgACC | PCu | | Voxel threshold of $p < .005$, cluster threshold > 10 adjacent voxels | N/A |
| Wang et al., 2017* | 23 | 25 | 38.74 (11.02) | Seed-based Analysis: PCC, MPFC, lLP, rLP, liTMP, riTMP, mdThal, lpCblm, rmCblm | no results | no results | Bonferroni correction: $p < .05/15$, network-wised; $p < .05/630$ ROI wised | 8 FE; 15 Recurrent |
| Zhu et al., 2017* | 31 | 32 | 20.5 (1.8) | Seed-based Analysis: 11 DMN ROIs | within dmPFC subsystem, between dmPFC and MTL subsystems | between core and dmPFC subsystems | FDR ($q < 0.05$) | All first-episode drug naïve |
| Knyazev et al. 2018* | 41 | 23 | 43.1 (13.8) | Seed-based Analysis: MPF and PCC | no results | no results | Permutation test ($p < 0.01$ to 0.001 voxel level, $p < 0.05$ cluster level) | N/A |

| | | | | | | | | |
|---|---|---|---|---|---|---|---|---|
| Evans et al., 2018* | 33 | 25 | 36 (10) | Seed-based Analysis: PCC | no results | no results | 3dClustSim: an initial threshold of $p < 0.05$ using a cluster size of > 120 | 6 (3) |

Abbreviations: fMRI, functional magnetic resonance imaging; MDD, major depressive disorder; FC, functional connectivity; ICA, independent component analysis; sgACC, subgenual anterior cingulate cortex; TFCE, threshold-free cluster enhancement; FDR, false discovery rate; GRF, Gaussian random field; DMN, default mode network; PCC, posterior cingulate cortex; mPFC, medial prefrontal cortex; dmPFC, dorsal medial prefrontal cortex; vmPFC, ventral medial prefrontal cortex; pgACC, perigenual anterior cingulate cortex; PCu, precuneus; AG, angular gyrus; LTG, lateral temporal gyrus; MTL, medial temporal lobe; rLTC, right lateral temporal cortex; pSTS, posterior superior temporal sulcus.

* Studies those were included in meta-analysis (Supplementary Figure S1).

† Summarize whether MDD patients are first-episode drug naïve or the number of previous episodes.

**Supplementary** Table S2. Existing or ongoing cohorts consisting MDD patients with functional brain imaging.

| Serial Number | Cohorts | Notes |
|---|---|---|
| 1 | Disease Imaging Data Archiving - Major Depressive Disorder Working Group (DIDA-MDD)(60) | A multi-site R-fMRI dataset with 709 MDD patients and 725 corresponding normal controls. All subjects were recruited from mainland China. Currently there is no data-sharing plan for this dataset. Recently a research (60) on the regional metrics abnormalities of MDD comparing to normal controls were published, but no result regarding alterations in FC among brain regions in MDD was reported. |
| 2 | The Netherlands Study of Depression and Anxiety (NESDA) (61) | A multi-site cohort study focusing on MDD as well as anxiety. Although the total sample size is large (N = 2850), only a small group of subjects (N = 200) were invited for brain fMRI scanning. A recent R-fMRI study from this cohort consist N = 140 (62). |
| 3 | BiDirect-Baseline (63) | A prospective study that aims at investigating the mutual relationship between depression and (subclinical) arteriosclerosis. Recently, the baseline wave of data collection has been done, including 999 patients with depression and 912 healthy controls (64). Some data from this cohort has been published. Two recent studies tried to discriminate MDD patients in this cohort (N = 180) from healthy controls (65) and divide them (N = 360) into subgroups (66) according to R-fMRI functional connectivity characteristics. |
| 4 | The Marburg-Münster Affective Disorders Cohort Study (MACS) (67) | An ongoing and a large cohort of sample (n = 2500) will be recruited. All participants will be scanned twice within 2 years. The fMRI data from recruited healthy participants (n = 444) (67), structural data from participants with unipolar depression (n = 58) and bipolar depression (n = 58) (68), and 74 MDD patients' task state fMRI data (69) have been published. |
| 5 | UK biobank (70) | A national and international health resource containing more than 4000 fMRI data from participants with depression. Although some structural MRI analysis have been done focusing on MDD patients from this cohort (71, 72), as far as we can see, no R-fMRI results of this MDD subsample have been published until now. |

| | | |
|---|---|---|
| 6 | The German National Cohort (GNC) (73) | A joint interdisciplinary endeavor aiming to investigate the causes for the development of major chronic diseases including MDD. Baseline data acquisition is ongoing, targeting N=30000 for brain imaging. No R-fMRI results of this MDD subsample have been published until now. |
| 7 | Multisite resting-state fMRI Initiative (PsyMRI) (74) | A multicenter-based utilization of already existing R-fMRI data, but are only accessible for members of the Consortium. The published studies were focusing on dementia, no R-fMRI results on MDD were published yet. |
| 8 | Decoded Neurofeedback (DecNef) Project (75) | A Japan based open access brain imaging sharing initiative. A recent R-fMRI study of this project only involved 93 MDD patients (76). |
| 9 | ENIGMA Major Depressive Disorder Working Group (77, 78) | The ENIGMA MDD Working group is an international collaboration currently including 35 research samples from 14 different countries worldwide, including brain scans from around >5000 MDD patients and >9000 controls. ENIGMA MDD's researches and projects are mainly focused on structural MRI. No R-fMRI studies are published until now. |

**Supplementary Table S3.** Samples of the REST-meta-MDD project, consortium sites, contributors, sample size, data acquisition parameters, and published studies based on the present cohorts.

| Serial Number | Sites (cohorts) | Principal investigators | Data organizer | N | | Scanner | Receive (coil) | TR (ms) | TE (ms) | Flip Angle (∘) | Thickness/ gap | Slice number | Time points | Voxel size | FOV | Published researches |
|---|---|---|---|---|---|---|---|---|---|---|---|---|---|---|---|---|
| | | | | MDD | NC | | | | | | | | | | | |
| 1 | National Clinical Research Center for Mental Disorders (Peking University Sixth Hospital) & Key Laboratory of Mental Health, Ministry of Health (Peking University) | Tian-Mei Si | Li Wang | 74 | 74 | Siemens Tim Trio 3T | 32 channel | 2000 | 30 | 90 | 4.0mm/0.8 mm | 30 | 210 | 3.28 × 3.28 × 4.80 | 210 × 210 | Wang et al 2013 (79)/2015 (80) |
| 2 | Department of Clinical Psychology, Suzhou Suzhou Psychiatric Hospital, The Affiliated Guangji Hospital of | Yan-Song Liu | Yan-Song Liu | 30 | 30 | Philips Achieva 3T | 8-channel | 2000 | 30 | 90 | 4.0mm/0 mm | 37 | 200 | 1.67 × 1.67 × 4.00 | 240 × 240 | Liu et al., 2017 (81) |

| | | | | | | | | | | | | | | | | |
|---|---|---|---|---|---|---|---|---|---|---|---|---|---|---|---|---|
| | Soochow University | | | | | | | | | | | | | | | |
| 3 | The Second Xiangya Hospital of Central South University | Shu-Qiao Yao / Xiang Wang | Chang Cheng | 27 | 37 | Siemens Magnetom Symphony scanner 1.5 T | 16 channel | 2000 | 40 | 90 | 5.0mm/1.25mm | 26 | 150 | 3.75 × 3.75 × 6.25 | 240 × 240 | Zhu et al., 2012 (24) |
| 4 | The Second Xiangya Hospital of Central South University | Wen-Bin Guo | Wen-Bin Guo | 24 | 24 | Siemens Skyra 3T | 32 channel | 2500 | 25 | 90 | 3.5mm/0 mm | 39 | 200 | 3.75 × 3.75 × 3.50 | 240 × 240 | Guo et al., 2014 (39)/2017(82) |
| 5 | Department of Psychiatry, Shanghai Jiao Tong University School of Medicine | Yi-Ru Fang / Dai-Hui Peng | Ru-Bai Zhou | 13 | 11 | GE Signa 3T | 32 channel | 3000 | 30 | 90 | 5.0mm/0 mm | 22 | 100 | 3.75 × 3.75 × 5.00 | 240 × 240 | Peng et al., 2014 (83)/2015 (84) |
| 6 | Department of Psychiatry, Shanghai Jiao Tong University School of Medicine | Yi-Ru Fang / Jun-Juan Zhu | Ru-Bai Zhou | 15 | 15 | Siemens Tim Trio 3T | 32 channel | 2000 | 30 | 70 | 4mm/0mm | 33 | 180 | 3.59 × 3.59 × 4.00 | 230 × 230 | Zhu et al., 2014 (85) |

| | | | | | | | | | | | | | | | | |
|---|---|---|---|---|---|---|---|---|---|---|---|---|---|---|---|---|
| 7 | Sir Run Run Shaw Hospital, Zhejiang University School of Medicine | Wei Chen | Jia-Shu Yao | 38 | 49 | GE discovery MR750 | 8 channel | 2000 | 30 | 90 | 3.2/0 | 37 | 184 | 2.29 × 2.29 × 3.20 | 220 × 220 | Shen et al., 2015 (86) |
| 8 | Department of Psychiatry, First Affiliated Hospital, China Medical University | Fei Wang | Jia Duan | 75 | 75 | GE Signa 3T | 8 channel | 2000 | 30 | 90 | 3.0mm/0 mm | 35 | 200 | 3.75 × 3.75 × 3.00 | 240 × 240 | Tang et al., 2013 (87) |
| 9 | The First Affiliated Hospital of Jinan University | Ying Wang | Guan-Mao Chen | 50 | 50 | GE Discovery MR750 3.0T | 8-channel | 2000 | 25 | 90 | 3.0/1.0 mm | 35 | 200 | 3.75 × 3.75 × 4.00 | 240 × 240 | N/A |
| 10 | First Hospital of Shanxi Medical University | Ke-Rang Zhang | Ai-Xia Zhang | 50 | 33 | Siemens Tim Trio 3T | 32 channel | 2000 | 30 | 90 | 3.0mm/1.52mm | 32 | 212 | 3.75 × 3.75 × 4.52 | 240 × 240 | Li et al., 2014 (88) |
| 11 | Department of Psychiatry, The First Affiliated Hospital of | Qing-Hua Luo / Hua-Qing | Hai-Tang Qiu | 32 | 29 | GE Signa 3T | 8 channel | 2000 | 30 | 90 | 5 mm | 33 | 200 | 3.75 × 3.75 × 5.00 | 240 × 240 | Du et al., 2016 (89) |

| | | | | | | | | | | | | | | | |
|---|---|---|---|---|---|---|---|---|---|---|---|---|---|---|---|
| | Chongqing Medical University | Meng | | | | | | | | | | | | | |
| 12 | Department of Psychiatry, The First Affiliated Hospital of Chongqing Medical University | Hua-Qing Meng / Qing-Hua Luo | Hai-Tang Qiu | 32 | 6 | GE Signa 3T | 8 channel | 2000 | 30 | 90 | 5 mm | 33 | 240 | 3.75 × 3.75 × 4.00 | 240 × 240 | N/A |
| 13 | The First Affiliated Hospital of Xi'an Jiaotong University, Xi'an Central Hospital | Jian Yang / Xiao-Ping Wu | Hong Zhang | 25 | 17 | GE Excite 1.5T | 16 channel | 2500 | 35 | 90 | 4mm/0 | 36 | 150 | 4.00 × 4.00 × 4.00 | 256 × 256 | Wu et al., 2016 (90) |
| 14 | The Second Xiangya Hospital of Central South University | Guang-Rong Xie | Xi-Long Cui | 64 | 32 | Siemens Tim Trio 3T | 32 channel | 2500 | 25 | 90 | 3.5/0 | 39 | 200 | 3.75 × 3.75 × 3.50 | 240 × 240 | Yang et al., 2017 (91) |
| 15 | Department of Psychosomatics and Psychiatry, Zhongda Hospital, School of Medicine, Southeast | Yong-Gui Yuan | Zheng-Hua Hou / Ying-ying Yin | 50 | 50 | Siemens Verio 3.0T MRI | 12 channel | 2000 | 25 | 90 | 4mm/0mm | 36 | 240 | 3.75 × 3.75 × 4.00 | 240 × 240 | Hou et al., 2018 (92)/2018 (93) |

| | | | | | | | | | | | | | | | | |
|---|---|---|---|---|---|---|---|---|---|---|---|---|---|---|---|---|
| | University | | | | | | | | | | | | | | | |
| 16 | Huaxi MR Research Center, West China Hospital of Sichuan University | Qi-Yong Gong / Kai-Ming Li | Kai-Ming Li | 31 | 31 | GE Signa 3T | 8 channel | 2000 | 30 | 90 | 5mm/0mm | 30 | 200 | 3.75 × 3.75 × 5.00 | 240 × 240 | Chen et al., 2017 (94) |
| 17 | Department of Psychiatry, The First Affiliated Hospital of Chongqing Medical University | Li Kuang | Lan Hu | 47 | 44 | GE Signa 3T | 8 channel | 2000 | 40 | 90 | 4.0mm/0 mm | 33 | 240 | 3.75 × 3.75 × 4.00 | 240 × 240 | Cao et al., 2016 (95) |
| 18 | Department of Radiology, The First Affiliated Hospital, College of Medicine, Zhejiang University | Hong Yang | Yu-Shu Shi / Hai-Yan Xie | 21 | 20 | Philips Achieva 3.0 T scanner (Philips Healthcare, Netherlands) | 8-channel SENSE head coil | 2000 | 35 | 90 | 5.0/1.0 mm | 24 | 200 | 1.67 × 1.67 × 6.00 | 240 × 240 | N/A |

| | | | | | | | | | | | | | | | | |
|---|---|---|---|---|---|---|---|---|---|---|---|---|---|---|---|---|
| 19 | Anhui Medical University | Kai Wang | Tong-Jian Bai | 51 | 36 | GE Signa 3T | 8 channel | 2000 | 22.5 | 30 | 4.0/0.6 mm | 33 | 240 | 3.44 × 3.44 × 4.60 | 220 × 220 | Wang et al., 2017 (96) |
| 20 | Faculty of Psychology, Southwest University | Jiang Qiu | Xin-Ran Wu | 282 | 251 | Siemens Tim Trio 3T | 12 channel | 2000 | 30 | 90 | 3.0mm/1.0 mm | 32 | 242 | 3.44 × 3.44 × 4.00 | 220 × 220 | Cheng et al., 2016 (97)/Ye et al., 2015 (98)/Luo et al., 2015 (99)/Xue et al., 2016 (100) |
| 21 | Beijing Anding Hospital, Capital Medical University | Chuan-Yue Wang | Qi-Jing Bo / Feng Li | 86 | 70 | Siemens Tim Trio 3T | 32 channel | 2000 | 30ms | 90 | 3.5mm/0.7 mm | 33 | 240 | 3.12 × 3.12 × 4.20 | 200 × 200 | Zheng et al., 2018 (101)/Jing et al., 2013 (102) |
| 22 | The Institute of Mental Health, Second Xiangya Hospital of Central South University | Zhe-Ning Liu | Yi-Cheng Long | 30 | 20 | Philips Gyroscan Achieva 3.0T | 32 channel | 2000 | 30 | 90 | 4.0mm/0 mm | 36 | 250 | 1.67 × 1.67 × 4.00 | 240 × 240 | N/A |

| | | | | | | | | | | | | | | | | |
|---|---|---|---|---|---|---|---|---|---|---|---|---|---|---|---|---|
| 23 | Mental Health Center, West China Hospital, Sichuan University | Tao Li | Yi-Ting Zhou | 32 | 30 | Philips Achieva 3.0T TX | 8 channal | 2000 | 30 | 90 | 4.0mm/0 mm | 38 | 240 | 3.75 × 3.75 × 4.00 | 240 × 240 | Yang et al., 2015 (103) |
| 24 | First Affiliated Hospital of Kunming Medical University | Xiu-Feng Xu / Yu-Qi Cheng | Chao-Jie Zou | 32 | 31 | GE Signa 1.5T | 8 channel | 2000 | 40 | 90 | 5/1mm | 24 | 160 | 3.75 × 3.75 × 6.00 | 240 × 240 | Cheng., et al. 2017 (104) |
| 25 | Department of Neurology, Affiliated ZhongDa Hospital of Southeast University | Zhi-Jun Zhang | Zhi-Jun Zhang | 89 | 63 | Siemens Verio 3T | 12 channel head coil | 2000 | 25 | 90 | 4.0mm/0 mm | 36 | 240 | 3.75 × 3.75 × 4.00 | 240 × 240 | Yuan et al., 2008 (105) |
| | Total | | | 1300 | 1128 | | | | | | | | | | | |

Abbreviations: MDD, major depressive disorder; NC, normal control.

**Supplementary Table S4.** Demographic characteristics for participants included in primary analysis.

| | Sites | Ranges across sites | | | | | | | | Grand mean and SD | | | | Group comparisons | |
|---|---|---|---|---|---|---|---|---|---|---|---|---|---|---|---|
| | N=25 | MDD (N=848) | | | | NC (N=794) | | | | MDD | | NC | | Mann-Whitney | |
| | n | min | max | mean | SD | min | max | mean | SD | mean | SD | mean | SD | Z | p |
| Age | 17 | 18-24 | 30-65 | 21.7-46.5 | 3.0-12.6 | 18-23 | 24-64 | 20.6-45.6 | 1.8-15.7 | 34.3 | 11.5 | 34.4 | 13 | 1.008 | 0.313 |
| Education | 17 | 3-9 | 15-21 | 9.7-14.2 | 1.5-4.5 | 5-12 | 15-23 | 9.9-15.9 | 1.6-4.8 | 12 | 3.4 | 13.6 | 3.4 | -10.2 | <0.001 |
| HAMD | 15 | 1-22 | 26-41 | 14.7-30.9 | 2.4-9.1 | | | | | 21.7 | 6.6 | | | | |
| Duration (month) | 15 | 0.2-9 | 12-480 | 5.3-90.1 | 4.2-102.5 | | | | | 38.4 | 60.6 | | | | |
| Episodes | 16 | 1-1 | 1-10 | 1-2.4 | 0-1.9 | | | | | 1.5 | 1.1 | | | | |
| | | | | | | | | | | | | | | Chi-Square | |
| | n | min | max | | | min | max | | | Sub n | % | Sub n | % | $X^2$ | p |
| Sex | | | | | | | | | | | | | | | |
| Male | 25 | 6 | 99 | | | 5 | 87 | | | 474 | 36.5 | 474 | 42.1 | 7.945 | 0.005 |
| Female | 25 | 5 | 183 | | | 1 | 164 | | | 826 | 63.5 | 653 | 57.9 | | |
| Subtype | | | | | | | | | | | | | | | |
| FEDN | 10 | 3 | 111 | | | | | | | 318 | 24.5 | | | | |
| Recurrent | 11 | 2 | 83 | | | | | | | 282 | 21.7 | | | | |

| | | | | | | | | | | | | |
|---|---|---|---|---|---|---|---|---|---|---|---|---|
| Unknown | 19 | 3 | 122 | | | | | | | | 700 | 53.8 |

**Supplementary Table S5.** Demographic characteristics for participants included in subgroup analysis

| | FEDN vs. NC | | | | | | recurrent vs. NC | | | | | | FEDN vs. recurrent | | | | | |
|---|---|---|---|---|---|---|---|---|---|---|---|---|---|---|---|---|---|---|
| | FEDN (N=232) | | NC (N=394) | | Mann-Whitney | | recurrent (N=189) | | NC (N=427) | | Mann-Whitney | | FEDN (N=119) | | recurrent (N=72) | | Mann-Whitney | |
| | mean | SD | mean | SD | Z | p | mean | SD | mean | SD | Z | p | mean | SD | mean | SD | Z | p |
| Age | 32.7 | 10.4 | 35.7 | 14.2 | -1.424 | 0.154 | 35.4 | 12.5 | 37.1 | 14.1 | -0.968 | 0.333 | 35.4 | 11.3 | 36.3 | 12.7 | -0.170 | 0.865 |
| Education | 12.2 | 3.4 | 13.6 | 3.6 | -5.702 | <0.001 | 11.7 | 3.2 | 13.4 | 3.8 | -6.162 | <0.001 | 11.5 | 3.4 | 12.0 | 3.5 | -1.104 | 0.270 |
| HAMD | 22.5 | 5.4 | | | | | 17.7 | 7.8 | | | | | 22.1 | 4.2 | 21.3 | 5.8 | 0.894 | 0.371 |
| Duration (month) | 17.7 | 30.8 | | | | | 92.7 | 86.1 | | | | | 27.0 | 39.5 | 88.7 | 80.1 | -6.419 | <0.001 |
| Episodes | | | | | | | 3.0 | 1.3 | | | | | | | 2.7 | 1.3 | | |
| | Male | Female | Male | Female | $X^2$ | p | Male | Female | Male | Female | $X^2$ | p | Male | Female | Male | Female | $X^2$ | p |
| Sex | 78 | 154 | 152 | 242 | 1.544 | 0.214 | 78 | 111 | 167 | 260 | 0.255 | 0.613 | 40 | 79 | 30 | 42 | 1.253 | 0.263 |

Abbreviations: FEDN, first episode drug naïve; NC, normal control.

**Supplementary Table S6.** Default mode network (DMN) within-network functional connectivity (FC) differences between 3 clinical subtypes.

| Contrasts | N | Age | Ratio of female | Mean FC within DMN (SD) | T (P) |
|---|---|---|---|---|---|
| CD+ vs. CD- | 92/97 | 32.26 (10.21)/32.71 (9.77) | 61.96%/56.70% | 0.29 (0.10)/0.27 (0.09) | 0.43 (0.67) |
| ANX+ vs. ANX- | 141/144 | 34.38 (10.46)/31.94 (9.92) | 65.96%/57.64% | 0.27 (0.25)/0.10 (0.08) | 0.48 (0.63) |
| NVSM+ vs. NVSM- | 121/129 | 33.55 (9.79)/30.38 (8.80) | 64.46%/58.91% | 0.26 (0.09)/0.26 (0.10) | -0.03 (0.98) |
| CD+ vs. ANX+ | 28/70 | 31.86 (11.07)/32.61 (10.18) | 53.57%/54.3% | 0.26 (0.10)/0.27 (0.09) | -0.94 (0.35) |
| CD+ vs. NVSM+ | 41/97 | 28.51 (8.63)/32.71 (9.77) | 46.34%/56.70% | 0.30 (0.12)/0.27 (0.09) | 1.46 (0.15) |
| ANX+ vs. NVSM+ | 58/130 | 30.88 (8.62)/31.14 (9.56) | 58.62%/56.92% | 0.28 (0.12)/0.24 (0.08) | 1.07 (0.29) |

Subtype definitions were based on Ahmed et al.'s mapping of HAMD scores to three National Institute of Mental Health Research-Domain-Criteria (RDoC) constructs: Core Depression (CD), Anxiety (ANX), and Neurovegetative Symptoms of Melancholia (NVSM) (15). CD+ was defined as those patients with a score of 3 or 4 on both HAMD items #1 and #7; ANX+ was defined by total score ⩾ 6 from items #9, #10, #11, and #15; NVSM+ was defined by a score of 1 or 2 on both items #6 and #12. Only participants who had HAMD item scores were included. When comparing two different clinical subtypes (e.g., CD+ vs. ANX+), subjects comorbid for both subtypes were excluded, thus subjects of one subtype in one contrast may differ from those in another contrast.

**Supplementary Table S7.** Verification results of default mode network (DMN) within-network functional connectivity (FC) in MDD with multiple alternative analysis strategies. Linear Mixed Effect (LME) model or meta-analytic model was utilized on different parcellations in different statistical comparisons (the effects of age, sex, education level, head motion and scanning site were controlled).

| | Dosenbach 160 functional ROIs (LME) | | Craddock 200 functional atlas (LME) | | Zalesky random 980 parcellations (LME) | | Dosenbach 160 functional ROIs (meta) | | Dosenbach 160 functional ROIs (LME & GSR) | | Dosenbach 160 functional ROIs (LME & Scrubbing) | |
|---|---|---|---|---|---|---|---|---|---|---|---|---|
| | *T* | *P* | *T* | *P* | *T* | *P* | *Z* | *P* | *T* | *P* | *T* | *P* |
| All MDDs vs. NCs (848 vs. 794) | -3.762 | 0.0002 | -2.638 | 0.008 | -3.179 | 0.002 | -4.057 | 0.00004 | -4.373 | 0.0001 | -3.818 | 0.0001 |
| FEDN MDDs vs. NCs (232 vs. 394) | -0.914 | 0.361 | -0.141 | 0.888 | -0.561 | 0.575 | -0.658 | 0.511 | -0.585 | 0.559 | -0.990 | 0.322 |
| Recurrent MDDs vs. NCs (189 vs. 427) | -3.737 | 0.0002 | -4.015 | 0.0001 | -3.356 | 0.0008 | -3.702 | 0.0002 | -4.382 | 0.0001 | -3.836 | 0.0001 |
| Recurrent MDDs vs. FEDN MDDs (72 vs. 119) | -2.676 | 0.008 | -3.064 | 0.003 | -3.284 | 0.001 | -1.732 | 0.083 | -0.974 | 0.331 | -2.527 | 0.012 |
| Long duration FEDN MDDs vs. Short duration FEDN MDDs (70 vs. 48) | 1.140 | 0.257 | 1.358 | 0.177 | 1.116 | 0.267 | 1.089 | 0.276 | 0.522 | 0.603 | 1.169 | 0.245 |
| Long duration MDDs vs. Short | 1.541 | 0.124 | 1.213 | 0.226 | 1.361 | 0.175 | 1.386 | 0.166 | 1.334 | 0.183 | 1.552 | 0.122 |

| | | | | | | | | | | | | |
|---|---|---|---|---|---|---|---|---|---|---|---|---|
| duration MDDs (186 vs. 112) | | | | | | | | | | | | |
| On medication MDDs vs. FEDN MDDs (115 vs. 97) | -2.629 | 0.009 | -2.359 | 0.019 | -2.293 | 0.023 | -2.568 | 0.010 | -1.891 | 0.060 | -2.504 | 0.013 |
| Correlation with HAMD in all MDDs (*N* = 734) | 1.591 | 0.112 | 1.576 | 0.116 | 1.181 | 0.238 | 0.754 | 0.451 | 0.448 | 0.654 | 1.765 | 0.078 |
| Correlation with HAMD in FEDN MDDs (N = 197 ) | -0.158 | 0.874 | 1.409 | 0.161 | 0.540 | 0.590 | -0.676 | 0.499 | -0.163 | 0.871 | -0.167 | 0.868 |
| Correlation with HAMD in recurrent MDDs (N = 126 ) | 2.167 | 0.032 | 1.424 | 0.157 | 1.264 | 0.209 | 1.304 | 0.192 | 1.741 | 0.084 | 2.446 | 0.016 |

Abbreviations: FEDN, First Episode Drug Na ïve; LME, Linear Mixed Effect; global signal regression; DMN, Default Mode Network.

**Supplementary Table S8.** *T* statistics of functional connectivity differences within- and between- 7 networks delineated by Yeo et al. (2011). Contrasts of All MDDs vs. NCs, FEDN MDDs vs. NCs, and Recurrent MDDs vs. NCs are listed in sequence separated by slashes.

| | VN | SMN | DAN | VAN | Subcortical | FPN | DMN |
|---|---|---|---|---|---|---|---|
| VN | -4.04*/-3.01*/-3.42* | | | | | | |
| SMN | -3.90*/-1.66/-4.14* | -4.00*/-1.81/-4.78* | | | | | |
| DAN | -3.86*/-1.55/-3.75* | -2.73*/-0.62/-3.67* | -2.08/-0.56/-2.12 | | | | |
| VAN | -1.67/0.05/-1.56 | -1.25/0.65/-1.53 | -0.59/1.41/-1.60 | -1.00/1.17/-1.99 | | | |
| Subcortical | -0.18/1.59/-0.32 | 0.39/2.23/-0.21 | 0.37/2.70/-0.56 | -0.05/1.95/-0.94 | -1.98/0.15/-2.22 | | |
| FPN | -0.71/-0.16/-1.16 | -1.83/-0.10/-1.99 | -1.61/-0.05/-1.74 | -0.38/0.05/-1.92 | -0.04/1.72/-1.05 | 0.34/0.48/-0.17 | |
| DMN | -0.78/0.06/-1.62 | -1.77/-0.95/-1.82 | -0.98/-0.90/-1.87 | 0.46/0.77/-1.13 | 0.63/1.87/-0.33 | 0.22/1.06/-1.14 | -3.76*/-0.91/-3.74* |

Abbreviations: VN, visual network; SMN: sensory-motor network; DAN: dorsal attention network; VAN: ventral attention network; Subcortical: subcortical ROIs; FPN: frontal parietal network; DMN: default mode network.

*: significant contrast after false discovery rate (FDR) correction. For the first contrast of comparing all 848 MDDs with 794 NCs, FDR correction was performed among 7 within-network and 21 between-network connections. For subgroup analyses, FDR correction was performed among the 6 abnormal connections found in the whole-group analysis.

**Supplementary Table S9.** *T* statistics of functional connectivity differences within- and between- 7 networks delineated by Yeo et al. (2011). Contrasts of Recurrent MDDs vs. FEDN MDDs, Long duration FEDN MDDs vs. Short duration FEDN MDDs, Long duration MDDs vs. Short duration MDDs, and On medication MDDs vs. FEDN MDDs are listed in sequence separated by slashes.

| | VN | SMN | DAN | VAN | Subcortical | FPN | DMN |
|---|---|---|---|---|---|---|---|
| VN | -0.75/0.65/0.86/-0.10 | | | | | | |
| SMN | -2.31*/-0.58/-0.24/-1.97 | -2.03/0.55/0.25/-2.17 | | | | | |
| DAN | -1.97/-0.19/0.26/-1.36 | -2.81*/0.89/-0.11/-2.52* | -2.62/0.87/-0.05/-2.42 | | | | |
| VAN | -1.23/-0.54/0.05/-0.18 | -1.87/1.18/-0.37/-1.27 | -1.79/0.36/-0.88/-0.93 | -2.50/0.7/-0.94/-1.53 | | | |
| Subcortical | -0.70/2.03/1.08/1.50 | -1.08/2.38/0.37/0.43 | -1.13/0.76/-0.35/0.25 | -1.63/1.71/-0.07/-0.84 | -1.78/0.36/-0.98/-1.22 | | |
| FPN | -1.18/0.99/0.76/0.21 | -1.91/0.71/-0.71/-0.85 | -1.54/0.68/-0.17/-0.86 | -1.39/0.89/-0.68/-0.36 | -1.16/-0.01/-0.05/-0.92 | -0.18/0.54/0.27/-0.05 | |
| DMN | -2.58/1.73/-0.18/-1.33 | -2.31/0.89/0.19/-1.56 | -2.17/0.36/-0.41/-1.29 | -2.36/0.04/-0.07/-1.71 | -1.87/0.04/0.58/-1.63 | -1.87/0.32/0.74/-2.43 | -2.68*/1.14/1.54/-2.63* |

Abbreviations: VN, visual network; SMN: sensory-motor network; DAN: dorsal attention network; VAN: ventral attention network; Subcortical: subcortical ROIs; FPN: frontal parietal network; DMN: default mode network.

*: significant contrast after FDR correction, performed among the 6 abnormal connections found in the whole-group analysis of Supplementary Table S8.

**Supplementary Table S10.** Correlation between within- and between- network functional connectivities and HAMD scores (presented in *T* values). Results calculated with all MDDs, FEDN MDDs and recurrent MDDs are listed in sequence separated by slashes.

| | VN | SMN | DAN | VAN | Subcortical | FPN | DMN |
|---|---|---|---|---|---|---|---|
| VN | -0.04/-0.11/0.15 | | | | | | |
| SMN | -0.02/0.06/1.28 | -1.31/-0.95/0.7 | | | | | |
| DAN | 0.46/0.65/0.88 | -0.21/-0.22/0.68 | 0.42/0.22/1.61 | | | | |
| VAN | 1.70/0.63/2.26 | 1.33/-0.51/1.60 | 1.42/0.63/0.97 | 0.33/-0.34/1.18 | | | |
| Subcortical | 2.11/0.93/1.46 | 1.74/0.21/1.81 | 2.15/0.6/2.11 | 1.43/-0.41/2.45 | -0.51/-2.55/1.81 | | |
| FPN | 1.81/0.99/0.77 | 1.54/1.23/0.57 | 1.22/1.00/1.41 | 0.99/0.62/0.83 | 1.75/0.51/1.92 | 0.03/-0.54/0.56 | |
| DMN | 1.82/0.52/0.86 | 1.42/1.14/0.93 | 1.54/1.33/1.09 | 1.73/0.86/1.73 | 1.69/0.11/2.06 | 1.04/-0.27/0.80 | 1.59/-0.16/2.17 |

Abbreviations: VN, visual network; SMN: sensory-motor network; DAN: dorsal attention network; VAN: ventral attention network; Subcortical: subcortical ROIs; FPN: frontal parietal network; DMN: default mode network.

*: significant correlation after FDR correction, performed among the 6 abnormal connections found in the whole-group analysis of Supplementary Table S8.

**Supplementary Table S11**. ROI pairs within DMN that are significantly different in functional connectivity between all MDDs and HCs. Both T values and corresponding p values are listed.

| Node 1 (MIN coordinates) | Node 2 (MIN coordinates) | T | P value |
|---|---|---|---|
| All MDDs vs. NCs | | | |
| vmPFC (6, 64, 3) | vmPFC (-6, 50, -1) | -2.99 | 0.0028 |
| vmPFC (6, 64, 3) | vmPFC (-11, 45, 17) | -3.92 | 0.0001 |
| vmPFC (6, 64, 3) | inf temporal (-61, -41, -2) | -3.76 | 0.0002 |
| vmPFC (6, 64, 3) | post cingulate (-5, -43, 25) | -3.81 | 0.0001 |
| vmPFC (6, 64, 3) | precuneus (9, -43, 25) | -2.97 | 0.0031 |
| vmPFC (6, 64, 3) | angular gyrus (-48, -63, 35) | -3.01 | 0.0027 |
| vmPFC (9, 51, 16) | inf temporal (-61, -41, -2) | -3.46 | 0.0006 |
| vmPFC (9, 51, 16) | precuneus (9, -43, 25) | -3.04 | 0.0024 |
| vmPFC (-6, 50, -1) | vmPFC (-11, 45, 17) | -3.13 | 0.0018 |
| vmPFC (-6, 50, -1) | ACC (9, 39, 20) | -3.47 | 0.0005 |
| vmPFC (-6, 50, -1) | inf temporal (-61, -41, -2) | -4.08 | <0.0001 |
| vmPFC (-6, 50, -1) | precuneus (-6, -56, 29) | -2.90 | 0.0038 |
| vmPFC (-11, 45, 17) | sup frontal (-16, 29, 54) | -3.43 | 0.0006 |
| vmPFC (8, 42, -5) | angular gyrus (51, -59, 34) | -3.50 | 0.0005 |
| vmPFC (8, 42, -5) | IPS (-36, -69, 40) | -2.95 | 0.0032 |
| sup frontal (23, 33, 47) | precuneus (9, -43, 25) | -3.46 | 0.0006 |
| sup frontal (-16, 29, 54) | inf temporal (-61, -41, -2) | -3.58 | 0.0003 |
| sup frontal (-16, 29, 54) | precuneus (9, -43, 25) | -3.29 | 0.0010 |
| sup frontal (-16, 29, 54) | post cingulate (-5, -52, 17) | -3.58 | 0.0004 |

| | | | |
|---|---|---|---|
| inf temporal (-59, -25, -15) | inf temporal (-61, -41, -2) | -3.20 | 0.0014 |
| inf temporal (-59, -25, -15) | precuneus (9, -43, 25) | -3.35 | 0.0008 |
| inf temporal (-61, -41, -2) | post cingulate (-5, -52, 17) | -2.89 | 0.0039 |
| post cingulate (-5, -43, 25) | precuneus (9, -43, 25) | -3.23 | 0.0013 |
| post cingulate (-5, -43, 25) | precuneus (5, -50, 33) | -3.39 | 0.0007 |
| post cingulate (-5, -43, 25) | post cingulate (-5, -52, 17) | -3.48 | 0.0005 |
| post cingulate (-5, -43, 25) | post cingulate (10, -55, 17) | -2.92 | 0.0035 |
| precuneus (9, -43, 25) | precuneus (5, -50, 33) | -3.54 | 0.0004 |
| precuneus (9, -43, 25) | post cingulate (-5, -52, 17) | -3.81 | 0.0001 |
| precuneus (9, -43, 25) | post cingulate (10, -55, 17) | -3.53 | 0.0004 |
| precuneus (9, -43, 25) | precuneus (-6, -56, 29) | -3.45 | 0.0006 |
| precuneus (9, -43, 25) | angular gyrus (51, -59, 34) | -2.92 | 0.0035 |
| precuneus (5, -50, 33) | post cingulate (10, -55, 17) | -3.48 | 0.0005 |
| precuneus (5, -50, 33) | precuneus (-6, -56, 29) | -3.62 | 0.0003 |
| precuneus (5, -50, 33) | post cingulate (-11, -58, 17) | -2.95 | 0.0032 |
| post cingulate (-5, -52, 17) | precuneus (-6, -56, 29) | -2.94 | 0.0034 |
| post cingulate (-5, -52, 17) | IPS (-36, -69, 40) | -3.23 | 0.0013 |
| post cingulate (10, -55, 17) | precuneus (-6, -56, 29) | -3.44 | 0.0006 |
| post cingulate (10, -55, 17) | angular gyrus (-48, -63, 35) | -3.06 | 0.0023 |
| post cingulate (10, -55, 17) | IPS (-36, -69, 40) | -3.4 | 0.0007 |
| angular gyrus (51, -59, 34) | angular gyrus (-48, -63, 35) | -2.96 | 0.0031 |
| angular gyrus (51, -59, 34) | IPS (-36, -69, 40) | -3.56 | 0.0004 |
| angular gyrus (-48, -63, 35) | IPS (-36, -69, 40) | -3.29 | 0.0010 |

Abbreviations: FEDN, first episode drug naïve; NC, normal control; vmPFC: ventral medial prefrontal cortex; inf: inferior; sup: superior; IPS: inferior parietal sulcus.

**Supplementary Table S12.** ROI pairs within DMN that are significantly different in functional connectivity between subgroups. Both T values and corresponding p values are listed.

| Node 1 (MIN coordinates) | Node 2 (MIN coordinates) | T | P value |
|---|---|---|---|
| Recurrent vs. NCs | | | |
| vmPFC (6, 64, 3) | post cingulate (1, -26, 31) | -3.14 | 0.0017 |
| vmPFC (6, 64, 3) | inf temporal (-61, -41, -2) | -3.7 | 0.0002 |
| vmPFC (6, 64, 3) | angular gyrus (51, -59, 34) | -3.52 | 0.0005 |
| vmPFC (6, 64, 3) | IPS (-36, -69, 40) | -4.59 | <0.0001 |
| mPFC (0, 51, 32) | post cingulate (-5, -43, 25) | -3.08 | 0.0022 |
| vmPFC (9, 51, 16) | inf temporal (-59, -25, -15) | -3.43 | 0.0006 |
| vmPFC (9, 51, 16) | angular gyrus (-48, -63, 35) | -3.51 | 0.0005 |
| vmPFC (-6, 50, -1) | inf temporal (-59, -25, -15) | -3.35 | 0.0009 |
| vmPFC (-6, 50, -1) | inf temporal (-61, -41, -2) | -3.93 | 0.0001 |
| vmPFC (-6, 50, -1) | angular gyrus (51, -59, 34) | -4.11 | <0.0001 |
| vmPFC (8, 42, -5) | inf temporal (-61, -41, -2) | -3.09 | 0.0021 |
| ACC (9, 39, 20) | vFC (51, 23, 8) | -3.15 | 0.0017 |
| ACC (9, 39, 20) | inf temporal (-61, -41, -2) | -3.43 | 0.0006 |
| ACC (9, 39, 20) | IPL (-53, -50, 39) | -3.22 | 0.0014 |
| sup frontal (-16, 29, 54) | inf temporal (-61, -41, -2) | -3.59 | 0.0004 |
| inf temporal (52, -15, -13) | angular gyrus (-48, -63, 35) | -3.32 | 0.0009 |
| precuneus (-3, -38, 45) | precuneus (5, -50, 33) | -3.23 | 0.0013 |
| post cingulate (-5, -43, 25) | post cingulate (-5, -52, 17) | -3.70 | 0.0002 |
| post cingulate (-5, -43, 25) | angular gyrus (51, -59, 34) | -3.82 | 0.0001 |

| | | | |
|---|---|---|---|
| sup temporal (42, -46, 21) | IPS (-36, -69, 40) | -3.29 | 0.0010 |
| post cingulate (-5, -52, 17) | angular gyrus (-48, -63, 35) | -3.29 | 0.0011 |
| post cingulate (10, -55, 17) | angular gyrus (-48, -63, 35) | -3.77 | 0.0002 |
| post cingulate (-11, -58, 17) | angular gyrus (-48, -63, 35) | -3.25 | 0.0012 |
| angular gyrus (51, -59, 34) | angular gyrus (-48, -63, 35) | -4.63 | <0.0001 |
| FEDN vs. NCs | | | |
| No significant difference | | | |
| Recurrent vs. FEDN | | | |
| mPFC (0, 51, 32) | ACC (9, 39, 20) | -4.23 | <0.0001 |
| ACC (9, 39, 20) | inf temporal (-61, -41, -2) | -3.74 | 0.0002 |
| angular gyrus (51, -59, 34) | angular gyrus (-48, -63, 35) | -4.03 | 0.0001 |
| First episode on medication vs. FEDN | | | |
| No significant difference | | | |

Abbreviations: FEDN, first episode drug na ïve; NC, normal control; vmPFC: ventral medial prefrontal cortex; inf: inferior; sup: superior; IPS: inferior parietal sulcus.

**Supplementary Table S13.** Functional connectivity between and within three sub-systems of DMN defined following Andrews-Hanna et al. (17): 1) core subsystem, 2) dorsal medial prefrontal cortex (dmPFC) subsystem, and 3) medial temporal lobe (MTL) subsystem. ROIs overlapped with the corresponding Yeo's 17 networks (18) were assigned to the subsystem as dissected by Andrews-Hanna et al. (2014) (17). Contrasts between all MDDs and NCs and sub-groups are listed.

| Contrast groups | | within core | between core-dmPFC | between core-MTL | within dmPFC | between dmPFC-MTL | within MTL |
|---|---|---|---|---|---|---|---|
| All MDDs vs. NCs | T | -4.842 | -2.939 | -3.011 | -2.707 | -1.458 | -2.136 |
| | p | <0.001 | 0.003 | 0.003 | 0.007 | 0.145 | 0.033 |
| FEDN MDDs vs. NCs | T | -1.003 | -0.513 | -0.929 | -0.487 | 0.007 | -1.232 |
| | p | 0.316 | 0.608 | 0.353 | 0.626 | 0.994 | 0.218 |
| Recurrent MDDs vs. NCs | T | -4.664 | -3.045 | -2.470 | -2.392 | -1.819 | -0.218 |
| | p | <0.001 | 0.002 | 0.014 | 0.017 | 0.069 | 0.827 |
| FEDN MDDs vs. Recurrent MDDs | T | 1.838 | 2.923 | 0.205 | 1.818 | 1.062 | 0.788 |
| | p | 0.068 | 0.004 | 0.838 | 0.071 | 0.290 | 0.432 |

**SUPPLEMENTARY FIGURES**

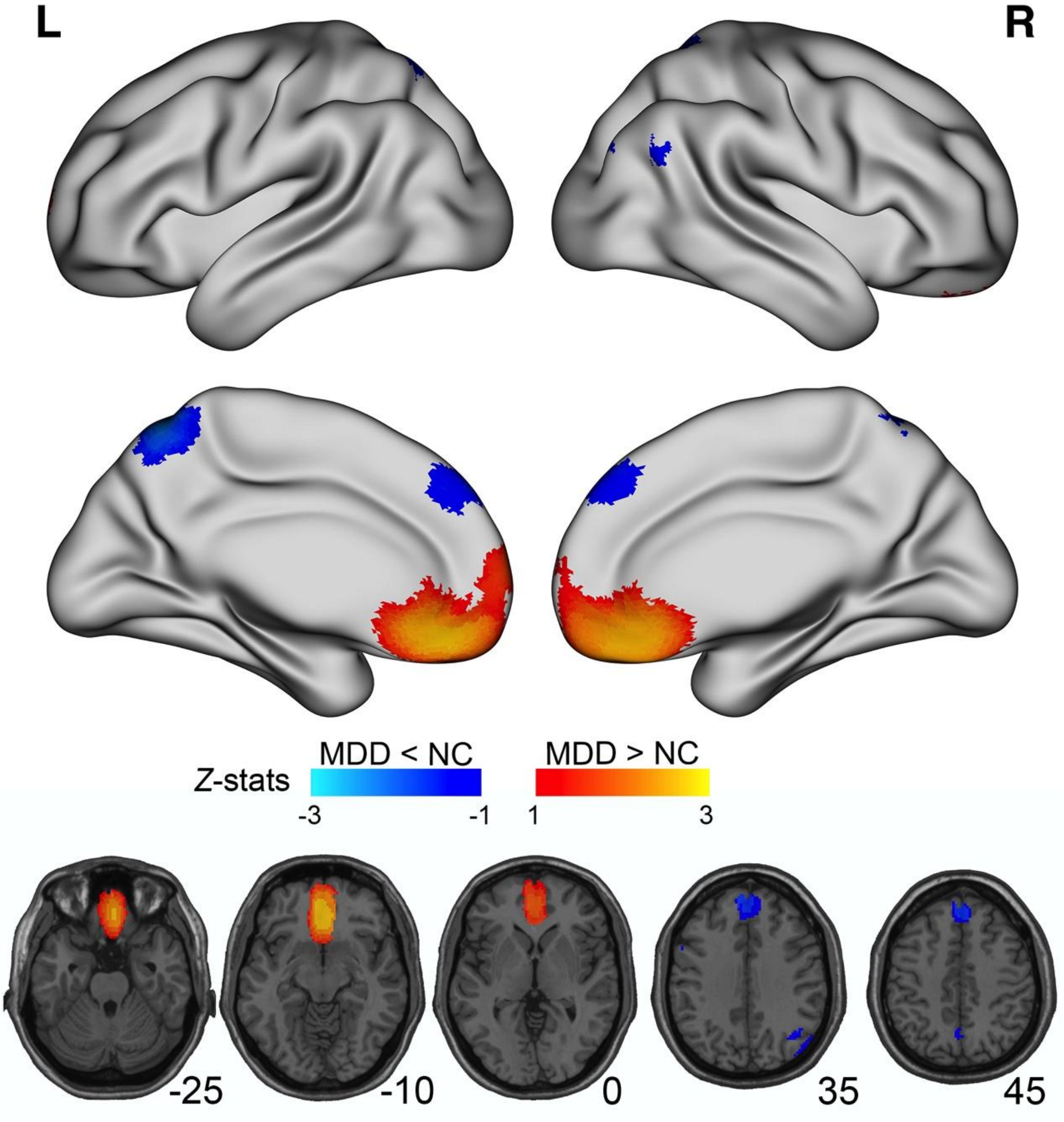


**Supplementary Figure S1.** Meta-analysis of existing literature on DMN FC in MDD. We further conducted a meta-analysis on the studies reviewed in Supplementary Table S1. Thirty-two whole brain voxel-wise studies which reported results with either Talairach coordinates or Montreal Neurologic Institute (MNI) coordinates were included in the meta-analysis. Signed Differential Mapping (SDM) (106) toolbox was used to identify spatially consistent differences of FC within MDDs' DMN compared to HCs. As described in detail in prior work (106), we first selected only peak coordinates (Talairach coordinates were transformed into MNI coordinates) which were statistically significant at the whole-brain level, with 5 studies reported no peaks (all the peaks and SDM table were shared through https://github.com/Chaogan-Yan/PaperScripts/tree/master/Yan_2018/Literature_meta_SDM). Secondly, for each individual study, an effect-size map of the MDD vs. HCs contrast differences of FCs was reconstructed by converting peak coordinate maximum t/p values into Hedges' effect sizes. Then an anisotropic Gaussian kernel was used to allocate greater effect sizes to voxels that were more highly correlated with the peaks. Finally, to account for the impact of sample size and intra-study and inter-study variability, we applied a random-effects model in which each study is weighted by the inverse of the sum of its variance plus the between-study variance as obtained by the DerSimonian-Laird estimator. The threshold was set as $p = 0.005$ (uncorrected) combined with $z > 1$, which was considered best balancing sensitivity and specificity while yielding results approximately equal to a corrected p value of 0.05 according to the original SDM paper (106). This meta-analysis showed increased orbitofrontal DMN FC and decreased dorsal medial prefrontal cortex (dmPFC) / posterior DMN FC in MDD.

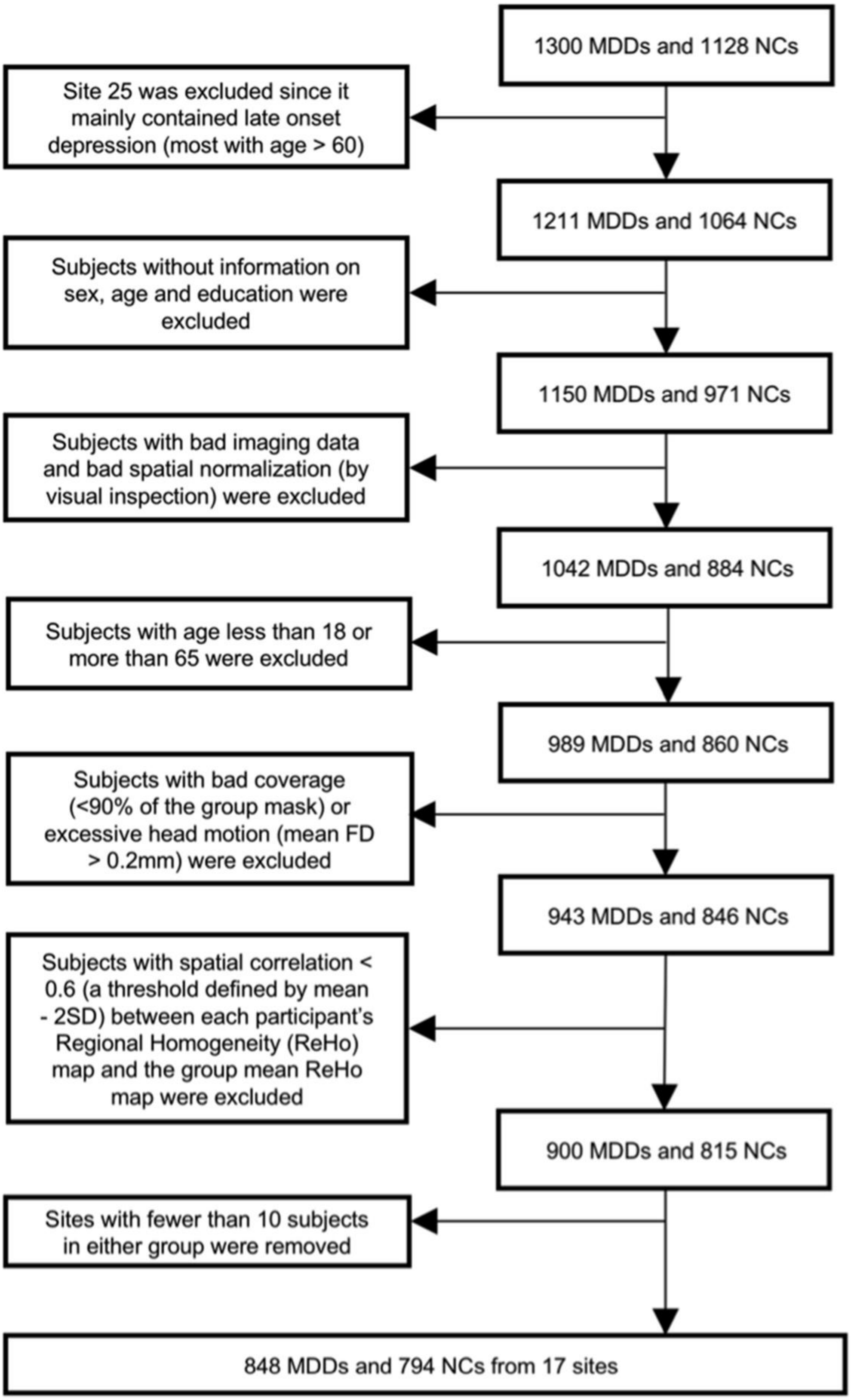


**Supplementary Figure S2.** Sample selection.

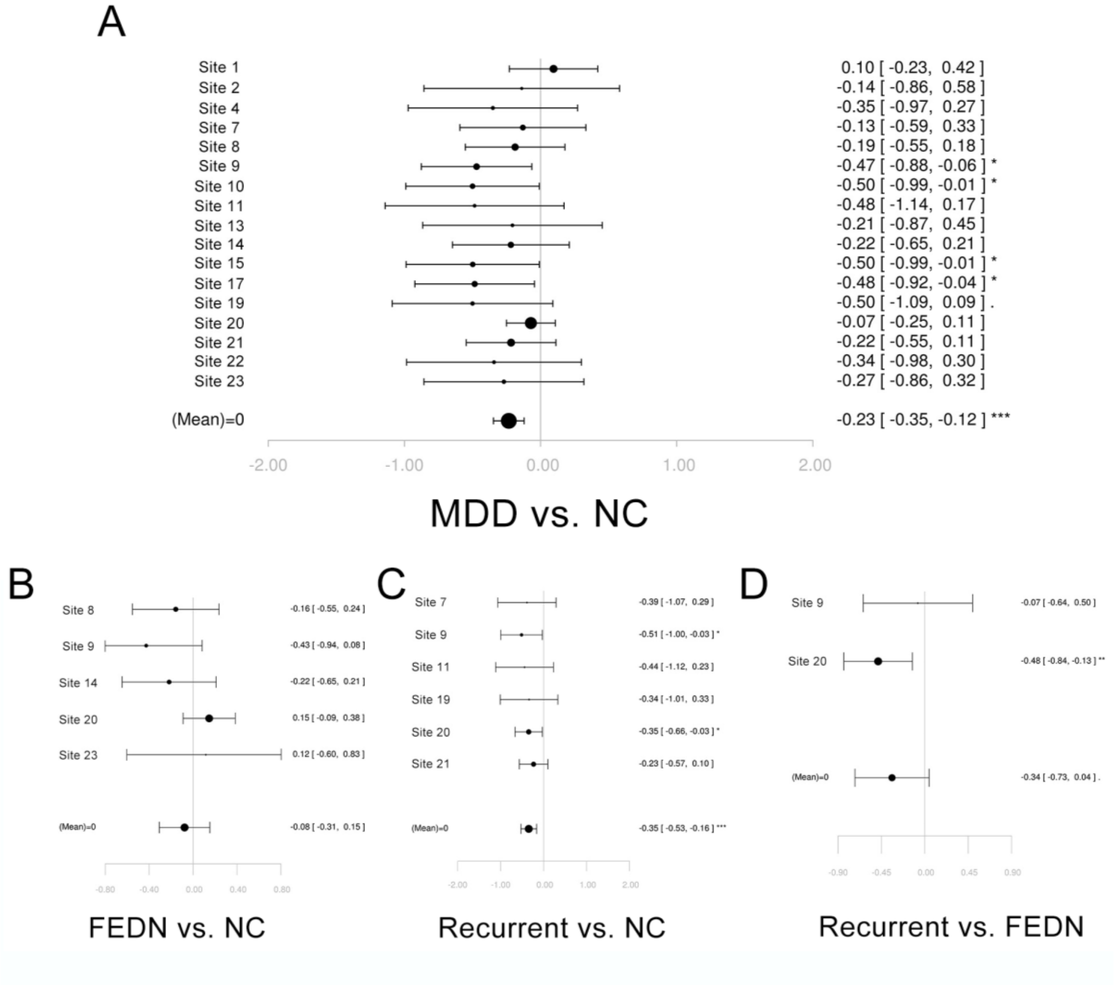


**Supplementary Figure S3.** Forest plots of effect size of each site generated by the meta-model in Reproducibility analysis: DMN within-network FC between MDD group and NC group (A), between first episode drug naïve (FEDN) MDD group and NC group (B), between recurrent MDD group and NC group (C), and between FEDN MDD group and recurrent MDD group (D). Of note, for each comparison, only sites with sample size larger than 10 in each group were included. The within-site T-values were calculated and converted into effect size, and then entered in a random effect meta-model using R package "metansue" (https://www.metansue.com/).

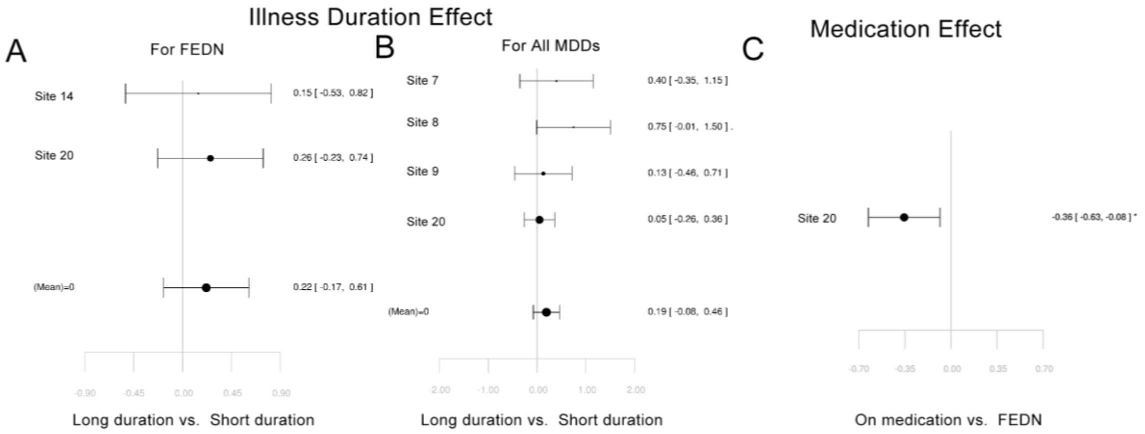


**Supplementary Figure S4.** Forest plots of effect size of each site generated by the meta-model in Reproducibility analysis: DMN within-network FC for first episode drug naïve (FEDN) MDDs with long vs. short illness duration (A), for pooled MDDs with long vs. short illness duration (B), and for first episode MDDs with vs. without medication usage (C). Of note, for each comparison, only sites with sample size larger than 10 in each group were included. The within-site T-values were calculated and converted into effect size, and then entered in a random effect meta-model using R package “metansue” (https://www.metansue.com/).

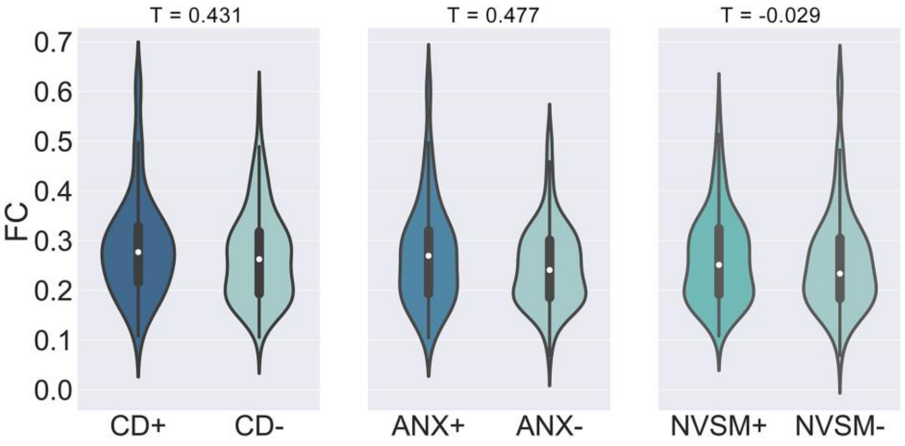


**Supplementary Figure S5.** Clinical Subtype effect on DMN within-network FC. The subtype definitions were based on Ahmed et al.'s mapping of the HAMD scale to the National Institute of Mental Health Research-Domain-Criteria (RDoC) constructs: Core Depression (CD), Anxiety (ANX), and Neurovegetative Symptoms of Melancholia (NVSM) (15). Please see details in Supplementary Table S6.

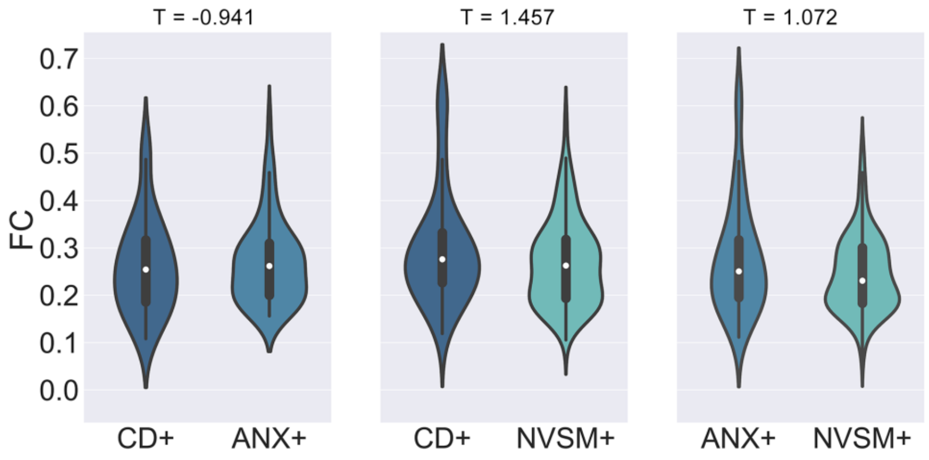


**Supplementary Figure S6.** Difference in DMN within-network FC between each pair of clinical subtypes while excluding subtype comorbidity. The subtype definitions were based on Ahmed et al.'s mapping of the HAMD scale to the National Institute of Mental Health Research-Domain-Criteria (RDoC) constructs: Core Depression (CD), Anxiety (ANX), and Neurovegetative Symptoms of Melancholia (NVSM) (15). Please see details in Supplementary Table S.

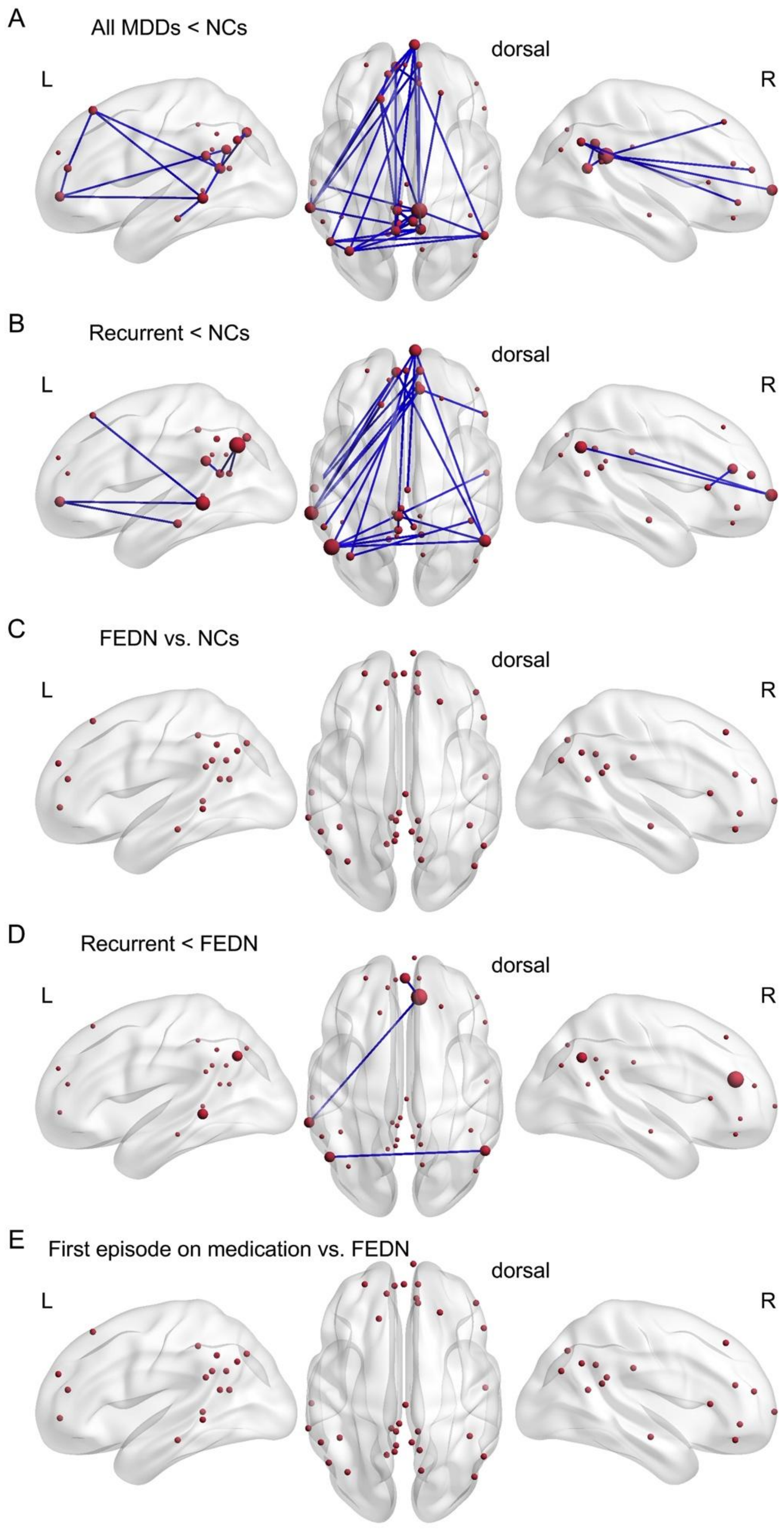


**Supplementary Figure S7.** Connection-wise decrease of DMN functional connectivity in MDD patients. The brain maps showed ROIs (red balls) defined within the DMN and connections (gray lines) between ROIs with significant decreased FC in each comparison, from the left-lateral view (left column), dorsal view (middle column), and right-lateral view (right column). The connection-wise comparisons of DMN FC were conducted between all MDDs vs. NCs (A), between recurrent MDDs vs. NCs (B), between first episode drug naïve (FEDN) MDDs vs. NCs (C), between recurrent vs. FEDN MDDs (D), and between first episode MDDs on medication with FEDN MDDs (E). Of note, all the tests were directional (two-tailed). L, left; R, right. The size of each red ball represented the number of its connections with significant group differences.

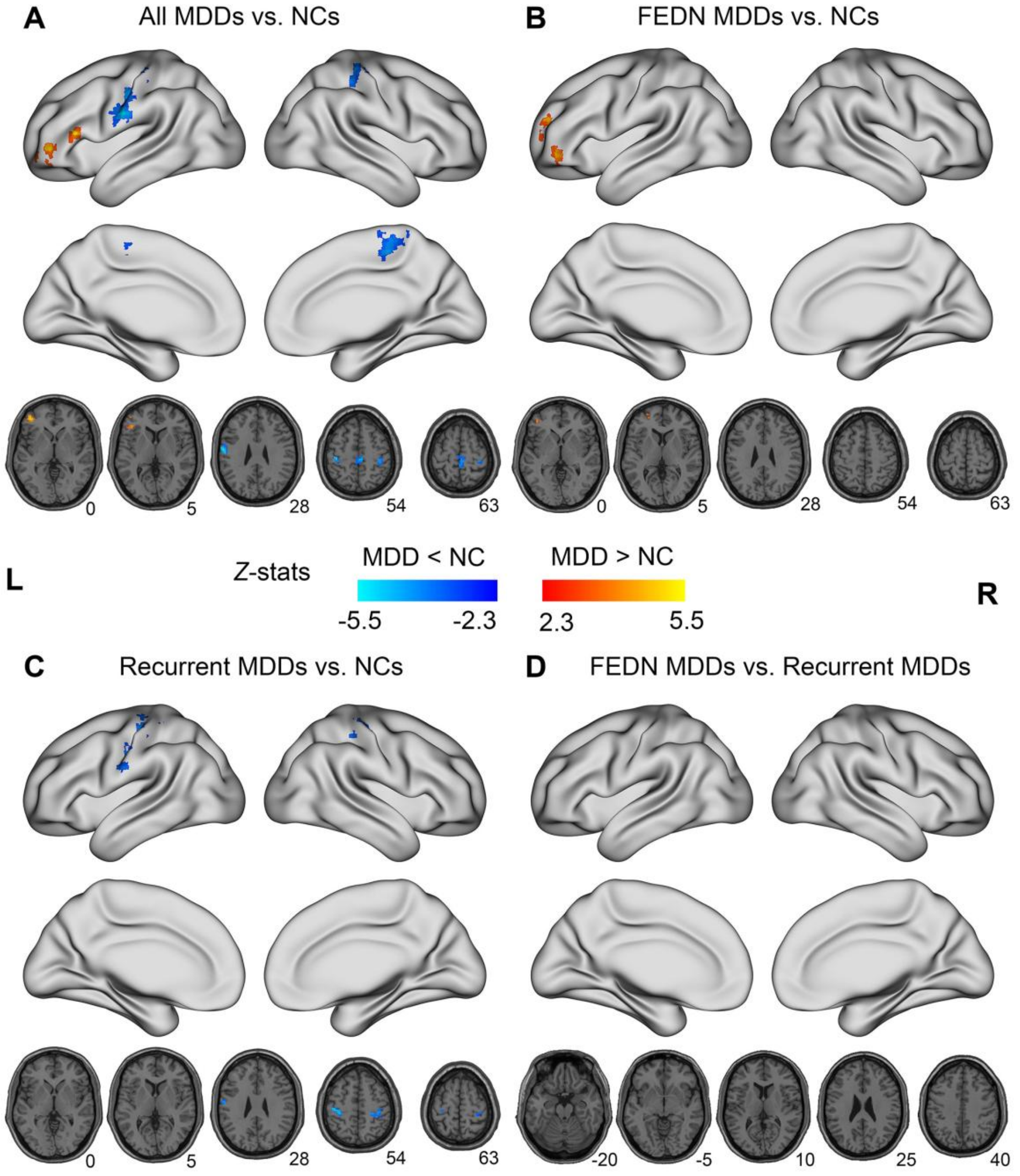


**Supplementary Figure S8**. The abnormalities of regional homogeneity (ReHo) in MDD patients. Significant group differences between (A) all individuals with MDD and NCs, (B) first episode drug naïve (FEDN) MDDs and NCs, (C) recurrent MDDs and NCs, (D) FEDN and recurrent MDDs are depicted. Gaussian random field (GRF) theory correction was employed to control family-wise error rates (voxel-level $p < 0.0005$; cluster-level $p < 0.025$ for each tail, two-tailed). L, Left hemisphere; R, right hemisphere.

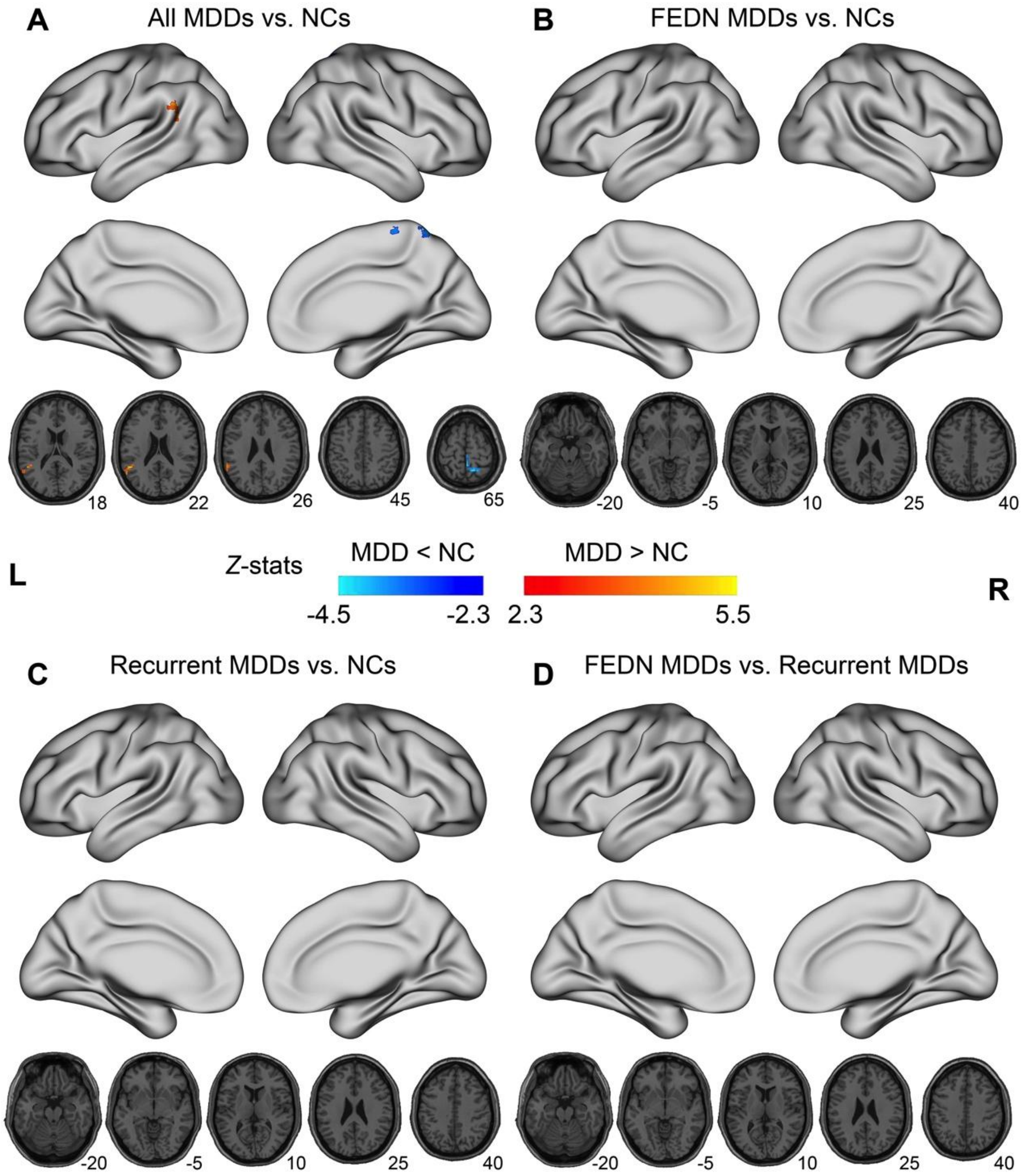


**Supplementary Figure S9**. The abnormalities of amplitude of low frequency fluctuations (ALFF) in MDD patients. Significant group differences between (A) all individuals with MDD and NCs, (B) first episode drug naïve (FEDN) MDDs and NCs, (C) recurrent MDDs and NCs, (D) FEDN and recurrent MDDs were depicted. Gaussian random field (GRF) theory correction was employed to control family-wise error rates (voxel-level $p < 0.0005$; cluster-level $p < 0.025$ for each tail, two-tailed). L, Left hemisphere; R, right hemisphere.

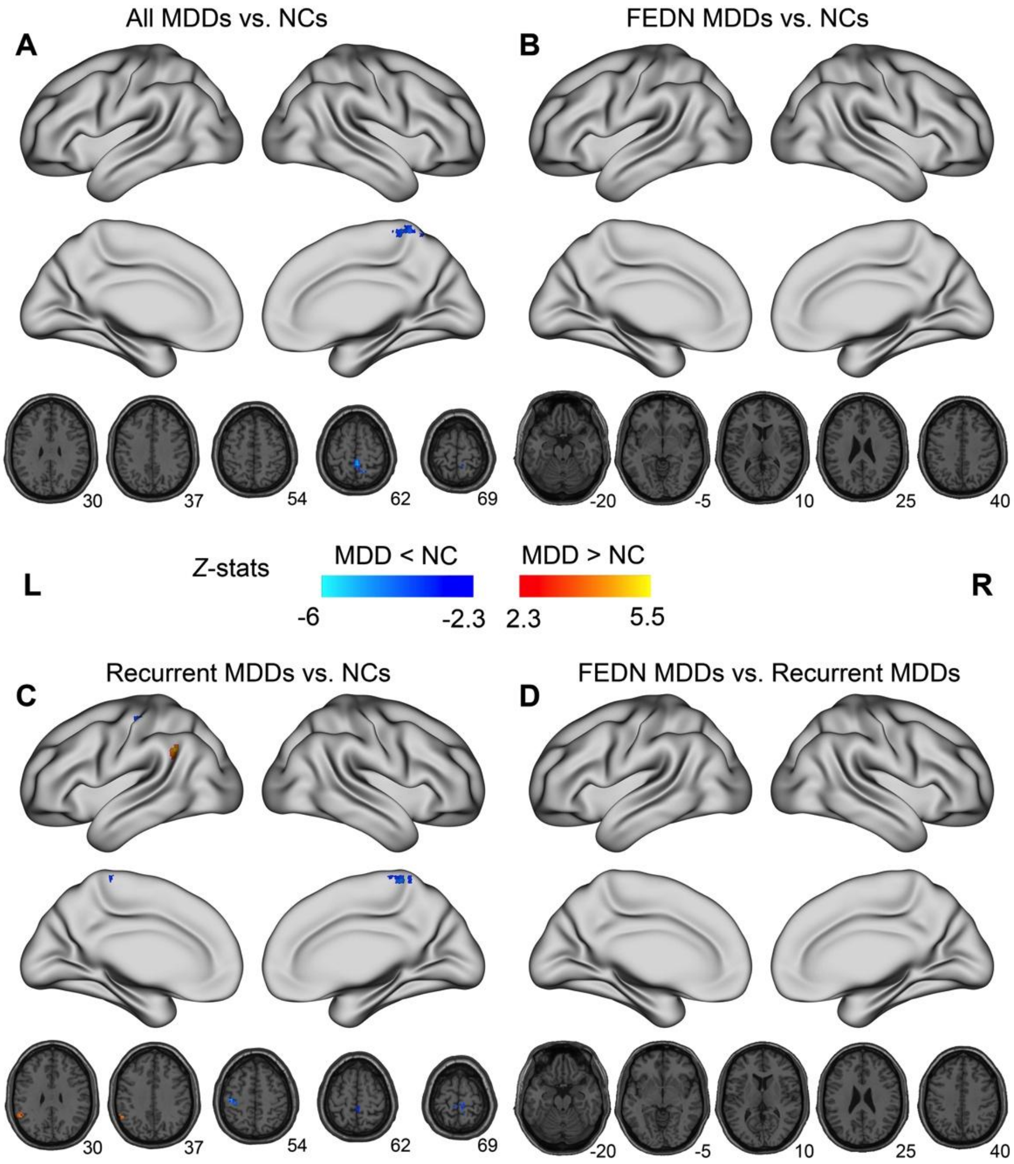


**Supplementary Figure S10**. The abnormalities of fractional ALFF (fALFF) in MDD patients. Significant group differences between (A) all individuals with MDD and NCs, (B) first episode drug naïve (FEDN) MDDs and NCs, (C) recurrent MDDs and NCs, (D) FEDN and recurrent MDDs were depicted. Gaussian random field (GRF) theory correction was employed to control family-wise error rates (voxel-level $p < 0.0005$; cluster-level $p < 0.025$ for each tail, two-tailed). L, Left hemisphere; R, right hemisphere.

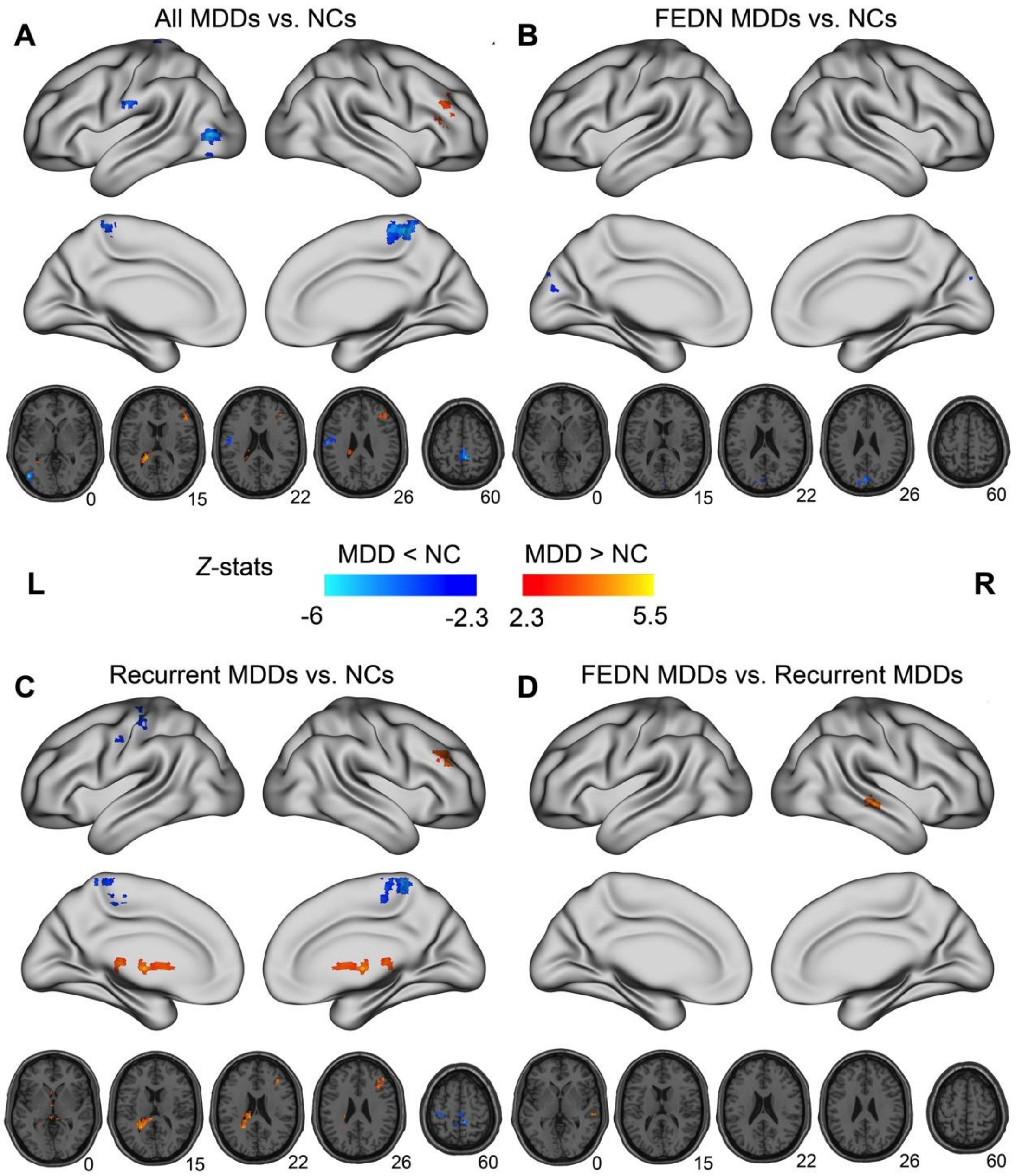


**Supplementary Figure S11**. The abnormalities of degree centrality (DC) in MDD patients. Significant group differences between (A) all individuals with MDD and NCs, (B) first episode drug naïve (FEDN) MDDs and NCs, (C) recurrent MDDs and NCs, (D) FEDN and recurrent MDDs were depicted. Gaussian random field (GRF) theory correction was employed to control family-wise error rates (voxel-level $p < 0.0005$; cluster-level $p < 0.025$ for each tail, two-tailed). L, Left hemisphere; R, right hemisphere.

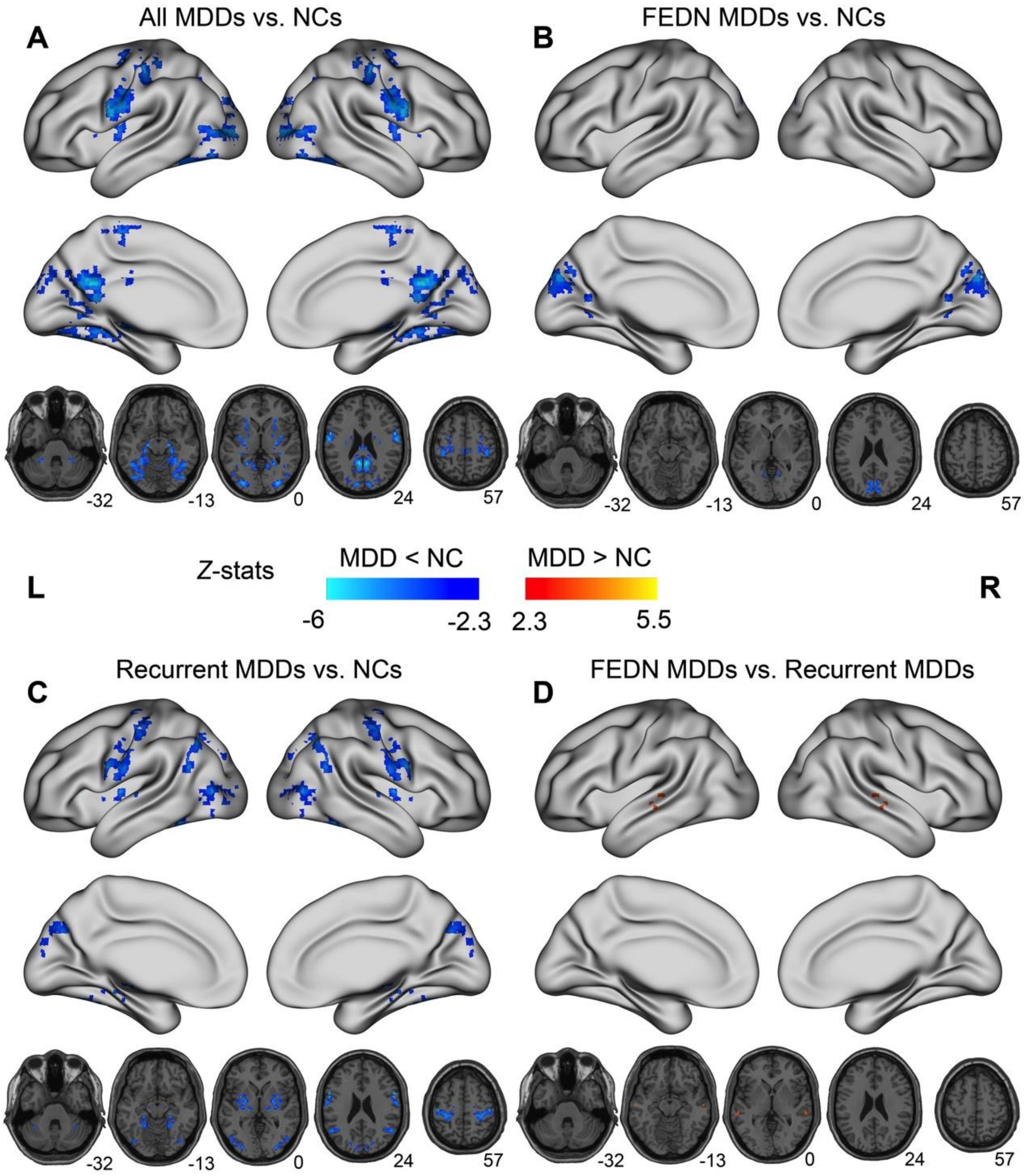


**Supplementary Figure S12**. The abnormalities of voxel-mirrored homotopic connectivity (VMHC) in MDD patients. Significant group differences between (A) all individuals with MDD and NCs, (B) first episode drug na ïve (FEDN) MDDs and NCs, (C) recurrent MDDs and NCs, (D) FEDN and recurrent MDDs were depicted. Gaussian random field (GRF) theory correction was employed to control family-wise error rates (voxel-level $p < 0.0005$; cluster-level $p < 0.025$ for each tail, two-tailed). L, Left hemisphere; R, right hemisphere.
